# Geometric Deep Learning Reveals Ligandable and Cryptic RNA Binding Small Molecule Pockets (SMARTPocket)

**DOI:** 10.64898/2026.06.18.732920

**Authors:** Riddhish H. Thakare, Amirhossein Taghavi, Jielei Wang, Jessica L. Childs-Disney, Chenglong Li, Matthew D. Disney, Yanjun Li

## Abstract

RNAs are important therapeutic targets, however identifying ligandable small-molecule binding pockets remains a major barrier to RNA-targeted drug discovery. Here, SMARTPocket, an atomic-level geometric deep learning framework for predicting RNA-small molecule binding pockets directly from three-dimensional structure is introduced. SMARTPocket represents RNA as full-atom point clouds and uses transfer learning from more than 110,000 protein binding interface structures to overcome the limited number of experimentally elucidated RNA-ligand complexes. Across four established single-chain benchmarks and three broader curated benchmarks, SMARTPocket consistently outperforms existing RNA pocket predictors and general biomolecular modeling approaches. The model generalizes to apo RNA structures when conformational changes are modest, identifies cryptic ligandable pockets, and recapitulates experimentally validated binding sites in the SARS-CoV-2 frameshifting element and an RNA aptamer evolved to bind small molecules. SMARTPocket-guided docking further improves near-native RNA-ligand pose recovery and computational efficiency compared with blind docking. These results establish SMARTPocket as a generalizable framework for structure-based identification of ligandable RNA pockets and for accelerating discovery of RNA-targeted small molecules.

## INTRODUCTION

RNAs are compelling therapeutic targets because they play central roles in regulating gene expression and numerous other cellular processes^1^. Advances in RNA biology have revealed that structured RNAs control diverse functions, including transcription, splicing, translation, RNA localization, and degradation, while dysregulation of these processes contributes to a wide range of human diseases^2^. As a result, small molecules that selectively recognize functional RNA structures have emerged as a promising therapeutic modality^3^. Central to these efforts is the identification of ligand-binding pockets within RNA, as these structural features provide the foundation for RNA-targeted drug discovery, structure-based ligand design^4^. Computational methods have emerged as cost-effective alternatives to experimental techniques, showing increasing promise for predicting RNA-small molecule binding pockets. However, existing approaches remain limited in accuracy and generalizability, as atomic features that define ligand recognition are missed^5–16^. Moreover, most approaches are designed for single-chain RNAs, even though functional pockets are often formed by multiple RNA chains or complex three-dimensional folds. These limitations are compounded by the relatively few RNA–small molecule complex structures available for training deep learning models. Some methods incorporate RNA dynamics, but these approaches are often computationally expensive and difficult to apply at scale^17^.

To address these limitations, we introduce **SMARTPocket** (**S**mall **M**olecule **A**pproaches to **R**NA **T**argeting **Pocket** Identification), an atomic geometric deep learning framework for RNA-small molecule binding pocket identification. SMARTPocket represents RNA structures as 3D atomic point clouds and utilizes a geometric transformer^18^ to learn hierarchical representations through attention mechanisms^19^, ultimately predicting nucleotide-level binding pocket probabilities. Because the model operates at the atomic level, it can use transfer learning from large biomolecular structure datasets, helping to overcome the scarcity of experimentally resolved RNA–ligand complexes.

Across multiple benchmark datasets, SMARTPocket shows improved or competitive performance compared to geometry-based methods (Rsite^5^, Rsite2^6^, RBind^7^, fpocketR^16^), feature-based machine learning methods (RNASite^9^, RLBind^11^, RNET^8^, ZHMolRestasite^10^), and data-driven deep learning methods based on sequence or structural information (RNABind^12^, GATRSite^13^, GeRNA-Bind^14^, SMARTBind^15^, AlphaFold3^20^). Moreover, it generalizes effectively to RNA targets with multiple chains and diverse structural motifs. Evaluation on apo (unbound) and holo (ligand-bound) conformations demonstrates that SMARTPocket can also identify potential ligand binding pockets in apo RNA structures, enabling binding site discovery even in the absence of known ligand or ligand bound states.

Our method was benchmarked against conventional docking approaches employed for the same pocket prediction task, and improvements in accuracy were observed, with markedly reduced target-to-target variability. Indeed, SMARTPocket can guide molecular docking to improve success rates and computational efficiency relative to blind docking. Case studies show that SMARTPocket recapitulates experimentally and chemically mapped RNA binding sites. It also identifies cryptic pockets^21^ that emerge after local conformational rearrangement and predicts ligandable regions in computationally modeled RNA structures, including FARFAR2 models^22^. Together, these results establish SMARTPocket as a generalizable framework for RNA pocket prediction and structure-guided discovery of RNA-targeted small molecules.

## RESULTS

### Model overview

SMARTPocket is a deep learning framework that uses a geometric transformer^18^ architecture to accurately predict RNA-small molecule binding pockets directly from RNA structural data (Fig 1. a). The model represents RNA structures as 3D atomic point clouds defined only by atom types and coordinates. Unlike many existing methods, SMARTPocket does not require handcrafted structural features, such as solvent-accessible surface area or predefined geometric descriptors. For each atom, SMARTPocket constructs a local geometric graph. Nodes represent atoms, and edges encode spatial relationships between neighboring atoms using interatomic distances and displacement vectors. A transformer based neural network then iteratively updates the node representations through attention mechanisms^19^, enabling effective information aggregation from the local neighbors. As the network becomes deeper, the number of nearest neighbors used for message passing progressively increases, allowing the model to capture broader structural context. Residual connections are incorporated throughout the network, mitigating vanishing gradient and over-smoothing^23^. An attention-pooling module then aggregates atom-level embeddings into nucleotide-level representations, and the final classifier assigns each nucleotide a probability of belonging to a small-molecule binding pocket.

**Fig. 1.**
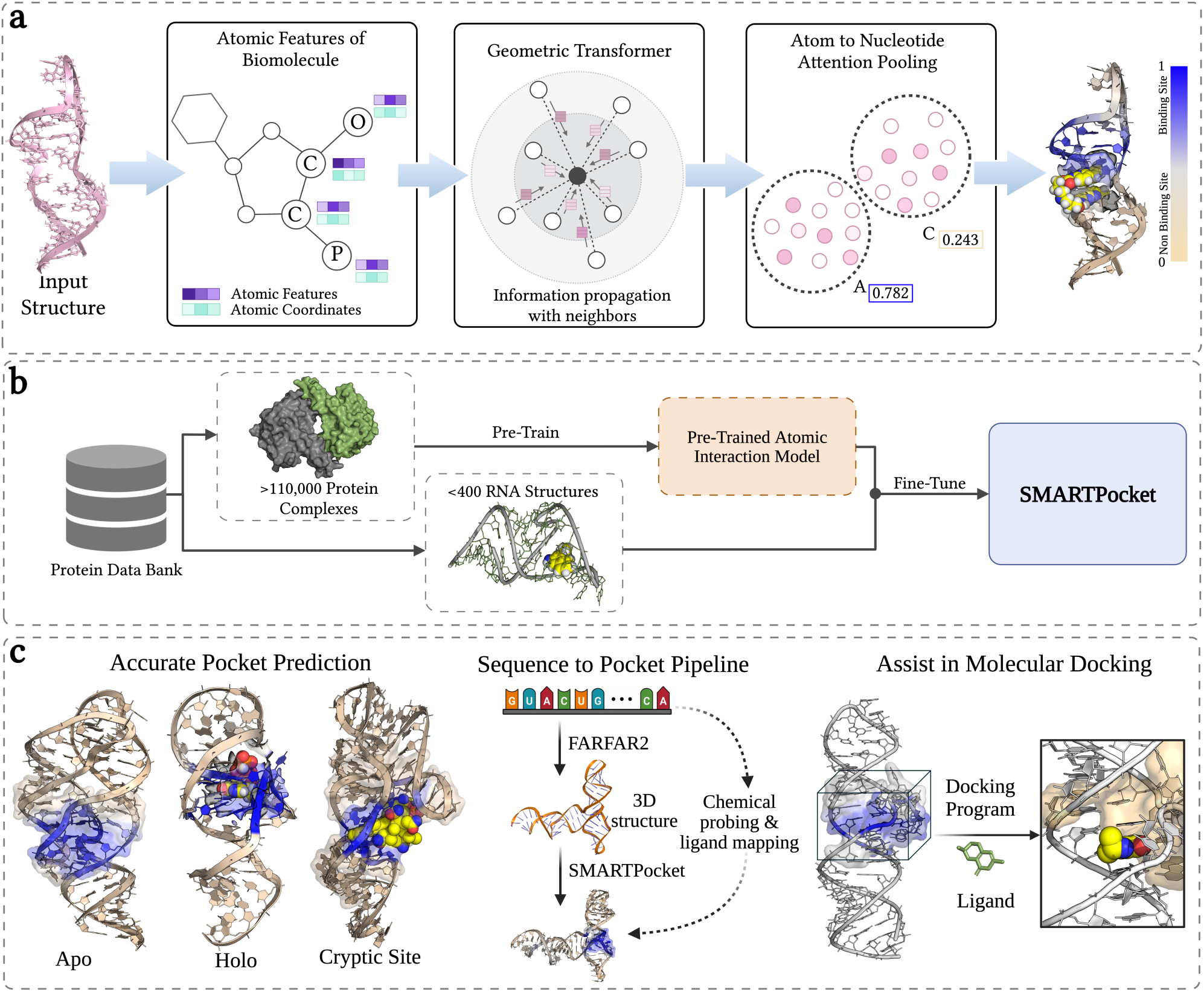
Overview of SMARTPocket framework for RNA-small molecule binding pocket prediction. **(a)** SMARTPocket represents RNA structures as atomic point clouds, from which graphs are constructed via nearest-neighbor connectivity. Geometric transformer layers propagate and update node and edge representations, and an attention-based pooling module aggregates atom-level representations into nucleotide-level embeddings, which are subsequently classified to yield binding site probabilities. **(b)** A transfer learning strategy was used to address RNA–small molecule data scarcity. The full-atom model was first pretrained on over 110,000 protein complexes and subsequently fine-tuned on two RNA training sets containing only 54 and 357 RNA structures, respectively, for the corresponding benchmarks. **(c)** SMARTPocket enables accurate RNA binding pocket identification across canonical and cryptic pockets, generalizes well to ligand-free (apo) RNA conformations, and recapitulates chemical probing–validated binding sites directly from computationally predicted RNA 3D structure. By defining focused ligandable regions, SMARTPocket can also guide molecular docking and improve the efficiency and accuracy of downstream pose prediction.

SMARTPocket uses translation-invariant and rotation-equivariant geometric operations, allowing RNA structures to be modeled in a physically consistent manner^18^. Translation invariance ensures that predictions do not depend on the absolute position of the RNA in the coordinate system. Rotation equivariance ensures that geometric representations transform consistently when the RNA is rotated. Together, these properties allow SMARTPocket to recognize binding pockets independent of the RNA’s position or orientation in space.

A major challenge for RNA pocket prediction is the limited number of experimentally elucidated RNA–small molecule complexes available for training^24, 25^; large and diverse datasets are required by deep learning models to learn molecular recognition patterns^26–29^. To address this challenge, a transfer learning strategy that pre-trained the model architecture on more than 110,000 protein interaction data following the PeSTo^18^ methodology was employed and subsequently fine-tuned on RNA-ligand structures (Fig. 1b). By operating at the atomic level, the model treats protein and RNA structures uniformly, enabling cross domain knowledge transfer and adaptation of molecular interface recognition priors to specific RNA-small molecule interactions. Detailed descriptions of the model architecture and training procedures are provided in the Methods section.

### Benchmarking SMARTPocket on single-chain RNA datasets

SMARTPocket was first evaluated on the widely used single-chain RNA benchmark datasets using the same training and testing protocol as the baseline methods. The training dataset (TR60) comprises 60 RNA-ligand structures and was partitioned using a 9:1 train-validation split to guide the model selection following the setting of RNASite^9^. Four single-chain RNA sets, including TE18^9^, RB9^7^, JL10^10^, and TL12^10^, served as the test sets. Detailed dataset information is provided in Methods section and Supplementary Table 1.

SMARTPocket was benchmarked against nine baseline methods: RSite^5^, RSite2^6^, RNASite^9^, RBind^7^, RLBind^11^, ZHmolReSTasite^10^, RNET^8^, RNABind^12^, and GATRSite^13^. Performance was evaluated using standard metrics, including area under the receiver operating characteristic curve (AUROC), Matthews correlation coefficient (MCC)^30^, F1 score, precision, and recall. A cumulative assessment across these complementary metrics is important as no single metric fully captures model performance, particularly for highly imbalanced residue-level binding site prediction tasks. AUROC evaluates the model’s overall ranking and discrimination capability across classification thresholds, whereas threshold-dependent metrics provide complementary perspectives: MCC measures balanced classification performance across positive and negative predictions, F1 score captures the trade-off between precision and recall, and Precision and Recall separately quantify the positive-prediction reliability and sensitivity, respectively. Together, these metrics provide a comprehensive and robust evaluation of predictive performance and model generalizability. Because all baseline methods used the same training and test sets, and some implementations are not publicly available, the performance metrics reported in the original or subsequent publications were used to ensure consistent comparison.

As summarized in Table 1, SMARTPocket demonstrated consistently strong performance across all test sets, often exceeding or matching the best-performing existing tools. On the TE18 test set, SMARTPocket achieved performance comparable to the leading GATRSite model^13^ (AUROC: 0.781 ± 0.007 vs. 0.780; MCC: 0.410 ± 0.026 vs. 0.320; mean ± s.d.). Notably, in this dataset, six of the 18 structures contain only ionic ligands, with no small molecules present. Because ionic ligands lack the three-dimensional pocket geometry that SMARTPocket is designed to recognize, their inclusion in the dataset likely reduced model performance observed on the full TE18 set. After excluding six such structures from dataset, SMARTPocket’s AUROC increased to 0.872 ± 0.010 (compared with 0.781 ± 0.007 on the full dataset), confirming that ligand composition rather than model capacity drives this difference.

**Table 1.** Performance comparison on the single-chain RNA benchmark datasets. Benchmarking SMARTPocket against nine baseline methods across four single-chain RNA test sets (TE18, RB9, JL10, and TL12). Performance is evaluated using AUROC, MCC, F1 score, precision, and recall. Baseline results for RLBind^11^, RNABind^12^, RNET^8^ and GATRSite^13^ results are taken from their original publications, and Rsite^5^, Rsite2^6^, RBind^7^, RNASite^9^, and ZHmolReSTasite^10^ are taken from the ZHmolReSTasite publication. Dashes indicate metrics that were not reported in the corresponding publication. SMARTPocket performance is reported as mean ± standard deviation over five random seeds. The best and second results are highlighted in bold and underlined.

| Datasets | Methods | AUC | MCC | F1 | Precision | Recall |
| --- | --- | --- | --- | --- | --- | --- |
| TE18 | Rsite | 0.474 | 0.071 | 0.351 | 0.449 | 0.288 |
|  | Rsite2 | 0.509 | 0.010 | 0.271 | 0.370 | 0.214 |
|  | RNASite | 0.776 | 0.253 | 0.378 | 0.675 | 0.263 |
|  | RBind | 0.559 | 0.187 | 0.273 | 0.655 | 0.173 |
|  | RLBind | 0.720 | 0.324 | <u>0.527</u> | 0.681 | 0.430 |
|  | ZHmolReSTasite | 0.709 | <u>0.327</u> | 0.498 | **0.729** | <u>0.379</u> |
|  | RNET | - | 0.307 | 0.473 | <u>0.701</u> | 0.357 |
|  | RNABind | 0.737 | - | - | - | - |
|  | GATRSite | <u>0.780</u> | 0.320 | 0.490 | 0.680 | 0.370 |
| | **SMARTPocket** | **0.781 ± 0.007** | **0.410 ± 0.026** | **0.668 ± 0.022** | 0.608 ± 0.013 | **0.742 ± 0.066** |
| RB9 | Rsite | 0.528 | 0.159 | 0.388 | 0.430 | 0.353 |
|  | Rsite2 | 0.575 | 0.072 | 0.251 | 0.382 | 0.187 |
|  | RNASite | 0.790 | 0.426 | 0.502 | 0.750 | 0.378 |
|  | RBind | 0.592 | 0.287 | 0.343 | 0.681 | 0.230 |
|  | RLBind | 0.851 | 0.546 | 0.660 | 0.750 | 0.590 |
|  | ZHmolReSTasite | 0.786 | 0.398 | 0.532 | 0.767 | 0.408 |
|  | RNET | - | 0.465 | 0.579 | **0.788** | 0.458 |
|  | RNABind | <u>0.865</u> | - | - | - | - |
|  | GATRSite | 0.850 | <u>0.570</u> | <u>0.680</u> | <u>0.770</u> | <u>0.610</u> |
| | **SMARTPocket** | **0.880 ± 0.016** | **0.574 ± 0.032** | **0.740 ± 0.026** | 0.665 ± 0.031 | **0.810 ± 0.108** |
| JL10 | Rsite | 0.477 | -0.046 | 0.234 | 0.295 | 0.194 |
|  | Rsite2 | 0.504 | 0.007 | 0.188 | 0.338 | 0.131 |
|  | RBind | 0.532 | 0.083 | 0.214 | 0.433 | 0.142 |
|  | ZHmolReSTasite | 0.592 | <u>0.211</u> | <u>0.384</u> | **0.549** | <u>0.296</u> |
|  | RNABind | <u>0.667</u> | - | - | - | - |
|  | **SMARTPocket** | **0.731 ± 0.014** | **0.330 ± 0.040** | **0.571 ± 0.052** | <u>0.508 ± 0.032</u> | **0.653 ± 0.128** |
| TL12 | ZHmolReSTasite | 0.704 | <u>0.440</u> | <u>0.606</u> | <u>0.740</u> | <u>0.514</u> |
|  | RNABind | <u>0.754</u> | - | - | - | - |
|  | **SMARTPocket** | **0.832 ± 0.020** | **0.545 ± 0.020** | **0.720 ± 0.072** | **0.804 ± 0.061** | **0.651 ± 0.088** |

More substantial gains were observed on other benchmarks. On the RB9 dataset, SMARTPocket achieved an AUROC of 0.880 ± 0.016, outperforming the second-best model, RNABind^12^ with an AUROC of 0.865. SMARTPocket’s advantage over RNABind likely stems from its atomic-level representation of RNA structures, which captures the precise atomic and geometric details governing ligand recognition. In contrast, RNABind has a coarse-grained nucleotide-level representation despite incorporating a pre-trained RNA large language model. Similarly, on the TL12 dataset, SMARTPocket achieved an AUROC of 0.832 ± 0.020 versus 0.754 for RNABind^12^. RB9 and TL12 datasets contain RNAs with moderate average chain sizes (51.0 and 46.9 nucleotides, respectively) which may explain the strong performance of SMARTPocket.

On the JL10 benchmark, SMARTPocket also delivered the highest performance (AUROC 0.731 ± 0.014 vs. 0.667 of RNABind^12^). Compared with the other test sets, the lower overall performance observed on the JL10 likely reflects the increased structural complexity of its constituent RNAs. Unlike simpler benchmark targets whose ligand-binding sites are often localized within hairpins, bulges, or internal loops, several JL10 structures contain multibranch junctions and higher-order tertiary architectures that generate binding pockets through long-range interactions between distant RNA segments. For example, the ligand-binding site in 9BUN is formed within a four-way junction where multiple helices and loops converge to create a tertiary pocket, while 7TZR contains a compact riboswitch architecture in which ligand recognition depends on the precise arrangement of multiple junction and helical elements. These pockets are therefore more complicated, making them inherently more difficult to identify computationally. Furthermore, four of the ten JL10 structures contain only ionic ligands, compounding the effect of structural complexity with the same ligand composition mismatch also observed in TE18. Excluding these four ion-only structures improves SMARTPocket’s AUROC to 0.766 ± 0.015 (compared with 0.731 ± 0.014 on the full dataset). Together, these results demonstrate that SMARTPocket provides strong and improved performance across diverse datasets.

### Benchmarking SMARTPocket on broader curated RNA–ligand datasets

To further evaluate the generalizability and robustness of our approach, a benchmark dataset derived from HARIBOSS^24^ (a comprehensive RNA-small molecule structural repository consisting of 868 structures retrieved from the Protein Data Bank (PDB)^31^) was curated. A multi-step filtration protocol was implemented to ensure high data quality and relevance for benchmarking RNA binding pocket prediction (See Methods). Wherein, ribosomal RNAs due to their structural complexity and RNA-ligand complexes where the ligand does not form a binding pocket (like end-stacking) were excluded, yielding a dataset of 447 RNA-small molecule complexes.

For data splitting, RNA targets were first partitioned based on pairwise structural similarity using hierarchical agglomerative clustering, following the protocol in RNAmigos2^32^ and RNAsmol^25^: an RMScore^33^ cutoff of 0.5 was applied such that RNAs with high structure similarity were assigned to the same cluster and therefore confined to a single split. This procedure yielded 25 clusters, which were divided into 357 training, 32 validation, and 58 test structures. To further minimize data leakage, an additional pocket-level similarity analysis was performed. Binding pockets were defined as all residues located within 4 Å of the bound ligand. Pockets in the test set were then compared against those in the training and validation sets using RMScore, and any test pocket exhibiting an RMScore greater than 0.5 to a pocket in either set was excluded reducing the final test set to 44 structures. An overview of the dataset filtering workflow is provided in Supplementary Fig. 7, and the PDB identifiers of all RNA structures are listed in Supplementary Table 2. Pairwise RMScore similarity heatmaps for RNA structures and binding pockets, confirming the absence of structural overlap between training, validation, and test sets, are shown in Supplementary Fig. 8.

Unlike previous benchmark datasets that focus solely on single-chain RNA, this dataset retains all RNA chains interacting with small molecules to ensure that binding pocket information is both accurate and complete. Further analysis of the HARIBOSS dataset reveals the high prevalence of multi-chain RNA structures, with 30% of the complexes consisting of 2–4 RNA chains (Supplementary Note 1). This is exemplified by the glmS ribozyme (PDB: 2H0Z, Supplementary Fig. 5), in which chains A and B jointly form the binding pocket for the competitive inhibitor Glc6P^34^, and by the guanine-binding site of the methyltransferase ribozyme MTR1 (PDB: 7Q7Z, Fig. 3b), which involves interactions among three RNA chains^35^.

### Evaluation on the HARIBOSS test dataset

Next, SMARTPocket was benchmarked against open source and web server-based baseline methods, including RNA-small molecule binding site predictors (RLBind^11^, RNABind^12^, SMARTBind^15^, fpocketR^16^, GerNABind^14^, and RNASite^9^) and the general purpose biomolecular structure prediction model AlphaFold3^20^. For RLBind and RNABind, each model was retrained on our training dataset five times using different random seed initializations, and the mean and standard deviation of the evaluation metrics were reported in Fig. 2. For fpocketR, GerNA-Bind, SMARTBind and AlphaFold3, inference using the provided software packages and model checkpoints were performed. For AlphaFold3, five co-folded structures were generated per sample using the RNA sequence and native ligand SMILES with different random seeds. For RNASite, binding site predictions were obtained via its web server. For methods producing per-nucleotide binding probability scores, a threshold of 0.5 was applied to classify each nucleotide as binding or non-binding. By contrast, fpocketR outputs binary predictions, whereas AlphaFold3 generates deterministic co-folded structures from which predicted binding nucleotides were extracted as those within 4 Å of the ligand pose in the predicted complex; both methods were therefore evaluated using threshold-independent metrics only.

**Fig. 2.**
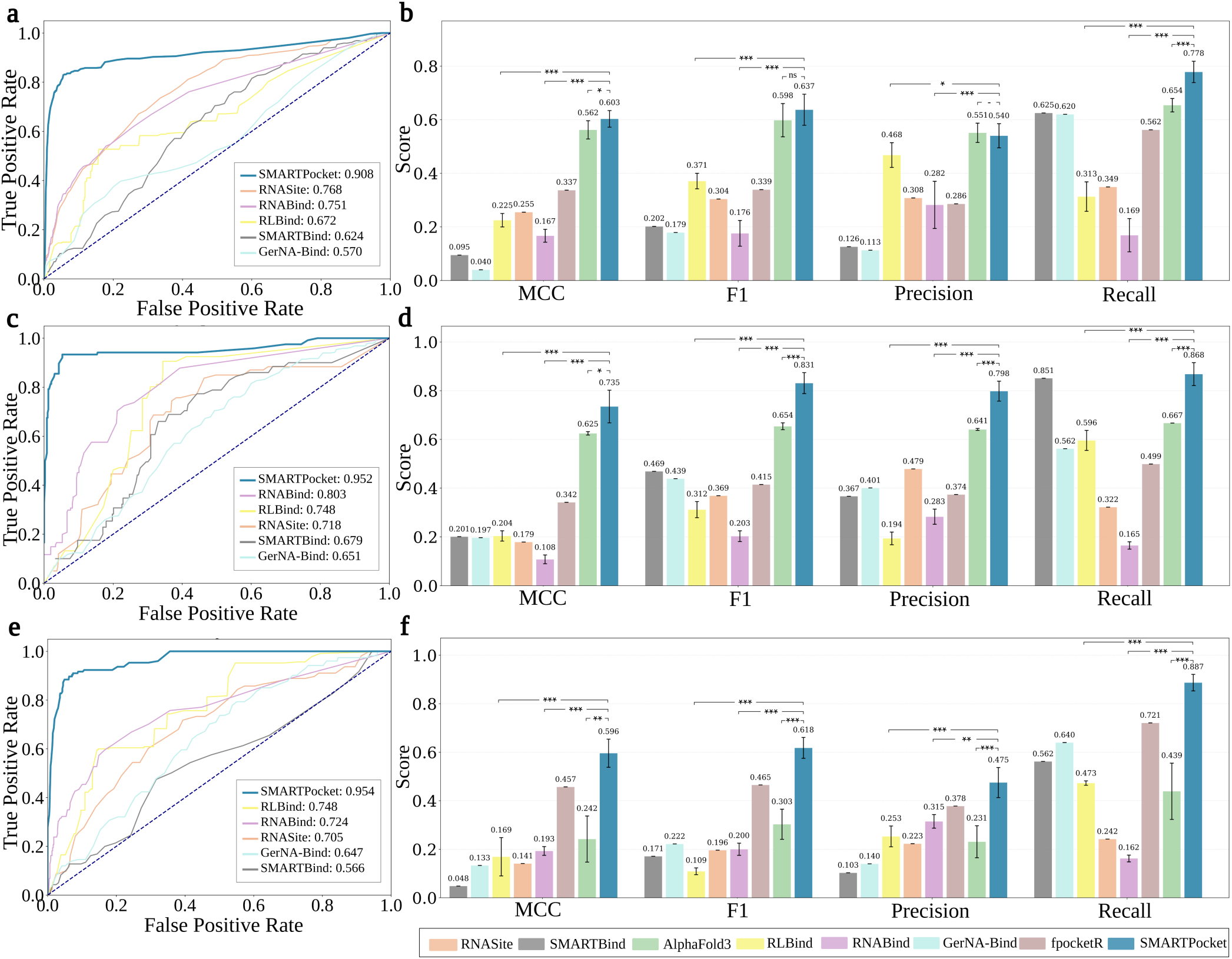
Performance comparison on the HARIBOSS, filtered Jiang, and the time-dependent test sets. **(a)** Receiver operating characteristic (ROC) curves on the HARIBOSS test set. SMARTPocket achieves the area under the ROC curve (AUROC = 0.908) among the six evaluated methods. The diagonal dashed line indicates random classification performance. **(b)** Detailed classification performance comparison of SMARTPocket and seven baseline methods on the HARIBOSS test set. Error bars represent the s.d. across runs; statistical significance was assessed using a one-tailed Wilcoxon rank-sum test (*P < 0.05, **P < 0.01, ***P < 0.001; n.s., not significant; –, not applicable). **(c)** ROC curves on the filtered Jiang test set, where SMARTPocket achieves the highest AUROC (0.952) among all evaluated methods on this independent benchmark. **(d)** Classification performance on the filtered Jiang test set, demonstrating that SMARTPocket maintains strong performance across MCC, F1, precision, and recall, with particularly high recall for binding site identification. Error bars represent the s.d. across runs. **(e)** ROC curves on the time-dependent test set, comprising RNA–small molecule complexes released after September 2021, where SMARTPocket achieves the highest AUROC (0.954) and demonstrates strong generalization to structurally novel targets absent from all methods’ training data. **(f)** Classification performance on the time-dependent test set, where SMARTPocket maintains consistently high performance across all metrics. Error bars represent the s.d. across runs.

As shown in Fig. 2a, SMARTPocket achieved consistently accurate binding site predictions, with a mean AUROC of 0.908 ± 0.016 (mean ± s.d.), compared to next-best baseline models, RNABind (0.751 ± 0.012) and RLBind (0.672 ± 0.022). SMARTPocket also attained the highest MCC of 0.603 ± 0.031, versus 0.562 ± 0.034 for the second-best model AlphaFold3 (Fig. 2b). Notably, AlphaFold3 was trained on biomolecule structure data through September 30, 2021^20^, and 36 of our 44 test samples fall within this timeframe, suggesting potential data overlap that may inflate its reported performance. Per-target performance of SMARTPocket is provided in Supplementary Table 3.

### Evaluation on external test sets

SMARTPocket was next evaluated on two external datasets. The first one was constructed from the Jiang Set^36^, which originally contains 800 noncovalent Nucleic Acid-ligand (NA-ligand) complexes. To maintain a strict external validation protocol, DNA-containing complexes and any RNA structures overlapping with or showing high RNA or pocket structural similarity (RMscore > 0.5) to any training and validation samples in our HARIBOSS dataset were removed. This resulted in six unique RNA-small molecule complexes for testing. The detailed curation process is described in Methods, and the resulting samples are listed in Supplementary Table 4.

On this dataset, SMARTPocket achieved strong predictive performance, with a mean AUROC of 0.930 ± 0.040 (mean ± s.d.), compared to 0.803 ± 0.025 for RNABind and 0.748 for RNASite (Fig. 2c). SMARTPocket also attained the highest MCC of 0.735 ± 0.067 and F1 score (0.831 ± 0.043), outperforming AlphaFold3 (0.625 ± 0.007 and 0.654 ± 0.014) and reflecting a strong balance between precision (0.798 ± 0.041) and recall (0.868 ± 0.047) as shown in Fig. 2d. While SMARTBind achieves comparable recall (0.851), its precision drops substantially to 0.367, indicating a high false positive rate in which a large proportion of predicted binding site residues are incorrectly assigned. This lack of specificity risks misdirecting downstream workflows by introducing spurious binding site residues. In contrast, SMARTPocket maintains high precision alongside high recall, making it considerably more reliable for downstream applications.

The second external benchmark is a time-dependent test set, consisting of RNA–small molecule complexes released in the PDB after September 30, 2021. To ensure strict independence from the training data, any samples with high RNA or pocket structural similarity to the HARIBOSS training and validation set were excluded (RMscore > 0.5; see Methods). The final dataset contains ten RNA–small molecule complexes, providing an independent temporal benchmark for model evaluation. Importantly, this temporal cutoff ensures no overlap with AlphaFold3’s training window. As shown in Fig. 2e-f, SMARTPocket achieved superior performance, with a mean AUROC of 0.954 ± 0.013 and an MCC of 0.596 ± 0.058 (mean ± s.d), surpassing all baselines including fpocketR (MCC 0.457), RNABind (MCC 0.193 ± 0.018), and AlphaFold3 (MCC 0.242 ± 0.095). The marked drop in AlphaFold3 performance likely reflects the absence of overlap with its training data. Notably, SMARTPocket achieved the highest precision (0.475 ± 0.062) alongside a high recall of 0.887 ± 0.034, a balance that is critical for accurate pocket localization while retaining sensitivity to novel targets. The full results for the ten cases are listed in Supplementary Table 5.

To better characterize the models’ ability to recover true binding site residues while limiting incorrect positive predictions, performance was also evaluated using the area under the precision–recall curve (AUPRC), which is particularly informative for the imbalanced binding site prediction task (see Supplementary Fig. 1). As shown in Supplementary Fig. 2a-c, SMARTPocket achieved the highest AUPRC values across all three datasets: 0.732 ± 0.033 on the HARIBOSS test set, 0.748 ± 0.041 on the filtered Jiang set, and 0.819 ± 0.052 on the time-dependent dataset. Notably, SMARTPocket maintained precision values exceeding 0.8 across a broad range of recall values, demonstrating reliable identification of binding site residues while limiting false-positive predictions. In contrast, competing methods exhibited more pronounced declines in precision over the same recall range, indicating reduced reliability of their predicted binding sites (Supplementary Fig. 2a-c).

### SMARTPocket enables accurate binding site prediction across diverse RNA 3D structural motifs

To assess model performance across diverse RNA structural motifs, RNA targets from the three curated test sets were aggregated and grouped according to RNA 3D Hub^37^ annotations for the structural regions surrounding the ligand-binding sites (Supplementary Table 8). As shown in Fig. 3a, SMARTPocket maintained strong performance across diverse RNA architectures, achieving the highest average AUROC across all six motif categories (0.926 ± 0.170) and outperforming state-of-the-art baselines (0.761 ± 0.211 for Alphafold3, 0.743 ± 0.203 for fPocketR, 0.741 ± 0.212 for RNABind, and 0.730 ± 0.209 for RNASite); differences were statistically significant (P < 0.001, one-tailed Wilcoxon rank-sum test). Similar trends were observed in the AUPRC radar plot (Supplementary Fig. 2d).

**Fig. 3.**
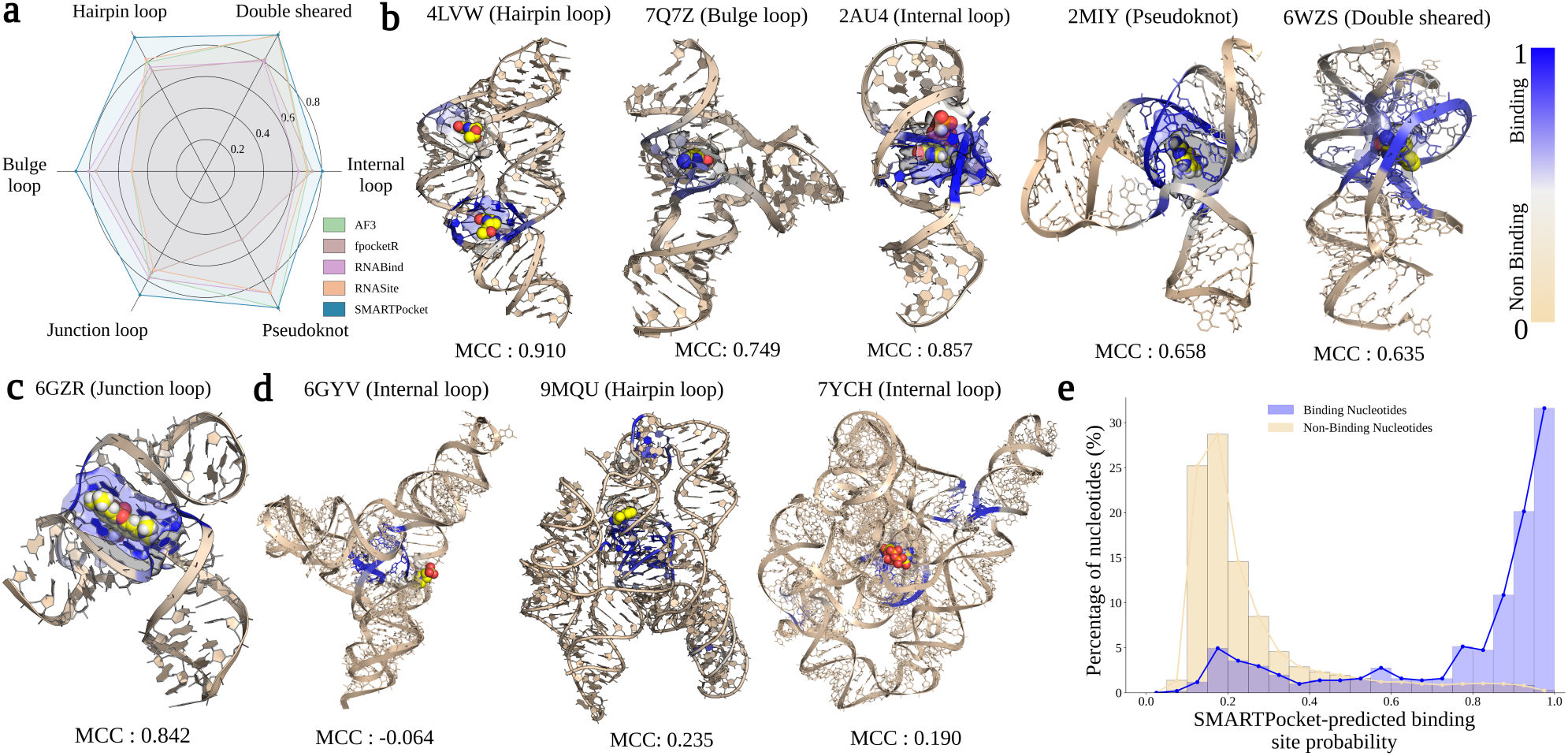
SMARTPocket accurately predicts binding pockets across diverse RNA structural motifs. **(a)** Radar plot comparing AUROC performance of SMARTPocket against four baseline methods across six structural motif classes, including hairpin, bulge, internal, and junction loops, double sheared and pseudoknot, evaluated on a combined dataset comprising the HARIBOSS, filtered Jiang, and time-dependent test sets. SMARTPocket consistently achieves superior AUROC across all motif classes. **(b, c)** SMARTPocket binding site predictions of representative examples with above diverse RNA motif classes. Ligands are rendered as spheres, while semi-transparent surfaces delineate the ground-truth binding sites. Predicted binding confidence is encoded using a tan-to-blue gradient (tan, non-binding; blue, binding) and MCC values are shown below each structure. **(d)** Representative cases in which SMARTPocket shows lower performance include 6GYV (internal loop, MCC = −0.064), 9MQU (hairpin loop, MCC = 0.235), and 7YCH (internal loop, MCC = 0.190). **(e)** Normalized distributions of SMARTPocket-predicted binding probabilities for true binding nucleotides (blue) and non-binding nucleotides (beige), showing well-separated distributions with binding nucleotides at high probability scores and non-binding nucleotides concentrated at low scores.

SMARTPocket predictions were further visualized on representative RNA structures. As shown in Fig. 3b-c, SMARTPocket assigned high binding site probabilities to the true ligand-binding regions in representative examples, including hairpin, bulge, internal and junction loops, as well as double-sheared motifs and pseudoknots. Notably, for the 4LVW case, which contains two distinct binding pockets, SMARTPocket accurately identified both sites and achieved an MCC of 0.910, highlighting its ability to detect multiple ligandable regions within a single RNA structure. For 7Q7Z, the binding pocket forms at the interface of multiple RNA chains, and SMARTPocket successfully localized it with an MCC of 0.749. Additional case studies are provided in Supplementary Fig. 5. Visual comparisons between SMARTPocket and AlphaFold3, the next-best performing model, across representative structures from the HARIBOSS and time-dependent test sets are provided in Supplementary Fig. 6.

A distribution of SMARTPocket performance across RNA sizes is provided in Supplementary Fig. 4. These results suggest that, despite the overall strong performance of SMARTPocket, precise prediction remains challenging for large RNAs or structures with complex conformations. Three representative cases illustrating the decline in SMARTPocket performance for larger RNA structures are shown in Fig. 3d. For the ribozyme 6GYV consisting of 192 nucleotides, SMARTPocket fails to accurately localize the binding pocket located within an internal loop, predicting a region away from the native ligand-binding position and yielding an MCC of −0.064. For the even larger structure 9MQU with 249 nucleotides, a drop in MCC to 0.235 was observed. In this structure, the predicted binding pocket is slightly displaced from the native ligand position, and an additional region was identified lacking an annotated ligand. Similarly, for 7YCH, a 404-nucleotide structure, SMARTPocket achieved an MCC of 0.190, where the predictions partially offset from the native binding site, covered a relatively broad area and included additional regions lacking annotated ligands. SMARTPocket’s difficulty to find binding sites in large RNAs likely reflects both the intrinsic complexity of these targets and the distribution of the training data; as shown in Supplementary Fig. 3, most training samples are concentrated in small- and medium-sized RNAs (<200 nt), resulting in limited model training exposure to larger or more structurally challenging cases.

The distribution of SMARTPocket predictions across the aggregated samples was also analyzed. Model outputs range from 0 to 1, representing the predicted probability that a nucleotide belongs to a true binding site, and nucleotide percentages were normalized separately for binding and non-binding labels. As shown in Fig. 3e, prediction scores for binding and non-binding nucleotides were clearly separated, with binding nucleotides predominantly concentrated in the high-confidence range of 0.8–1.0. This separation further indicates that top-ranked SMARTPocket predictions are highly enriched for true binding pockets and are therefore more reliable for guiding the downstream applications.

### Generalization of SMARTPocket to apo RNA structures

While SMARTPocket demonstrates strong performance across all benchmark datasets, it is important to recognize that these datasets consist exclusively of ligand-bound (holo) RNA structures. In practice, RNA molecules are highly flexible and frequently undergo conformational rearrangements upon ligand binding^38, 39^. As a result, the holo structures used for benchmarking may differ from the apo structures that exist prior to ligand engagement. In practical RNA drug discovery settings, the goal is often to identify ligand-binding pockets in RNAs for which no ligand binders are known, meaning only ligand-free (apo) structures are available. This presents a significant challenge, as ligand binding can stabilize conformations that are often only transiently populated or not readily apparent in the apo state^40, 41^.

Building on the success of SMARTPocket with holo structures, the generalizability of the model to apo RNA conformations was investigated. Using a set of six apo–holo RNA pairs curated in the SHAMAN^17^ and DRLiPS^42^ study, binding site predictions on both structural states were assessed. Overlap of either the apo or holo structure with the HARIBOSS training and validation sets was prevented by removing RNA targets with RMScore-based structural similarity greater than 0.6 to structures in these sets. Performance was quantified by computing the AUROC of both structures and Spearman correlation (Fig. 4a) between apo and holo predictions. Higher apo AUROC and correlation values indicate stronger model generalization to apo structures.

**Fig. 4.**
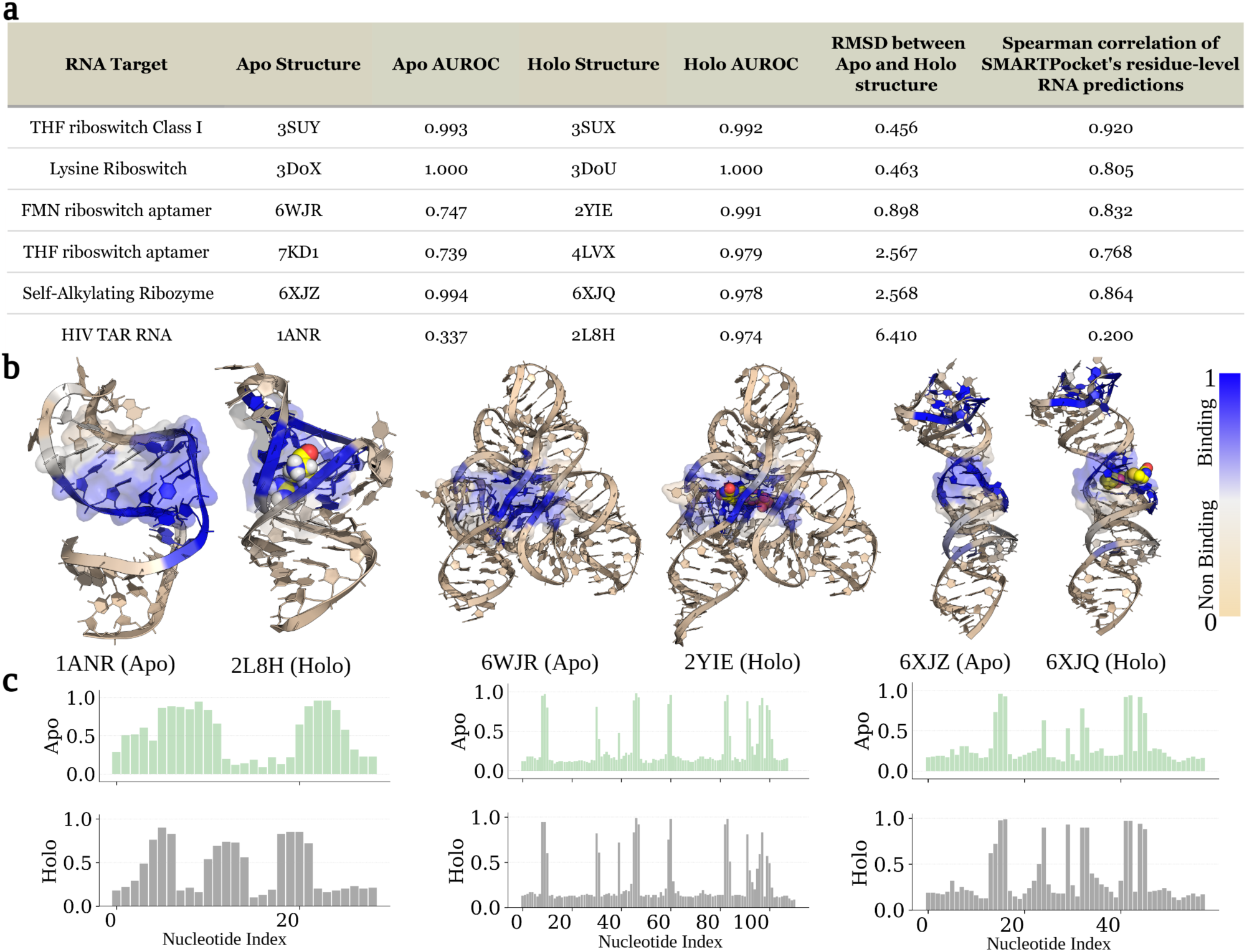
SMARTPocket performance across apo-holo RNA conformational changes. **(a)** SMARTPocket performance across six RNA targets with paired apo–holo conformations. The table summarizes AUROC values for both apo and holo states, root mean square deviation (RMSD) values between the two conformations (calculated in PyMOL^58^), and Spearman correlation between SMARTPocket residue-level binding predictions between apo and holo states. **(b)** SMARTPocket binding site predictions of three representative RNA targets with paired apo (left) and holo (right) conformations. Ligands in holo structures are rendered as spheres. Predicted binding confidence is encoded by a tan-to-blue gradient (tan, non-binding; blue, binding). **(c)** Histograms of predicted binding probability (y-axis) as a function of nucleotide position (x-axis) for apo (green, top) and holo (gray, bottom) states. Concordant peaks reflect consistent pocket identification across conformations, while discordances highlight regions subject to conformational reorganization upon ligand binding.

As shown in Fig. 4a, SMARTPocket achieved consistently high performance on holo structures, with AUROC values exceeding 0.97 across all six RNA targets examined, confirming its ability to accurately identify RNA binding pockets in ligand-bound structures. On apo structures, SMARTPocket maintained comparable accuracy and high prediction consistency with the corresponding holo conformations when structural differences were small, whereas performance declined as apo–holo structural divergence increased. For cases with similar apo and holo structures (RMSD < 1.0 Å), including THF riboswitch class I, the lysine riboswitch, and FMN riboswitch aptamer, prediction correlations remained consistently high (Spearman > 0.8), indicating reliable performance under limited conformational change. At intermediate divergence (RMSD ≈ 2.6 Å), performance was more variable: SMARTPocket retained strong accuracy for the self-alkylating ribozyme (apo AUROC = 0.994, Spearman = 0.864) but showed reduced performance on the THF riboswitch aptamer (apo AUROC = 0.739, Spearman = 0.768). By contrast, cases involving larger conformational rearrangements, such as HIV TAR RNA (RMSD = 6.410 Å), showed substantially reduced correlation (0.200) and poor apo AUROC prediction (0.337). Together, these results indicate that SMARTPocket reliably identifies ligand-binding pockets from apo structures when RNA conformation changes are limited, while performance decreases in cases where ligand binding induces major structural reorganization. Fig. 4b presents representative case studies of three apo-holo pairs, with corresponding nucleotide-wise binding predictions shown in Fig. 4c. Detailed per-target performance across additional metrics is provided in Supplementary Table 9.

### SMARTPocket identifies cryptic and Chem-CLIP validated RNA binding sites

To evaluate SMARTPocket’s ability to detect both canonical and noncanonical ligand-binding pockets directly from RNA structure, two structurally and functionally distinct systems were examined: a recently reported cryptic RNA binding site^21^ that emerges through local nucleotide displacement, and the SARS-CoV-2 frameshifting element (FSE)^43^, a viral regulatory RNA with experimentally validated small-molecule binding pockets.

Cobalamin riboswitches can access a cryptic binding pocket when A20 is displaced from its stacked position between G19 and A68, creating additional space for bulky ligands^21, 44^. In native or hydroxocobalamin-bound structures, A20 remains stacked, restricting access to this region (PDB 4FRG^45^; Fig. 5a, left). When bound to larger derivatives, A20 shifts toward the major groove, revealing a cavity of >340 Å³ that supports extended aromatic substituents. SMARTPocket reflected this structural plasticity such as in the pre-displacement state, ligandability scores were lower for G19 and A68 (0.465 and 0.247, respectively) and higher for A20 (0.856), consistent with the intact stacked configuration. Upon A20 displacement, predicted ligandability increased across G19, A68, and A20 (G19: 0.833, A68: 0.822, and A20: 0.893), matching the expanded accessibility of the cryptic site (PDB 9E5M; Fig. 5a, right). Notably, SMARTPocket assigned detectable ligandability to the region before displacement, indicating the tractability of the cryptic pocket. Detailed performance across the cobalamin riboswitch structures is provided in Supplementary Table 6.

**Fig. 5.**
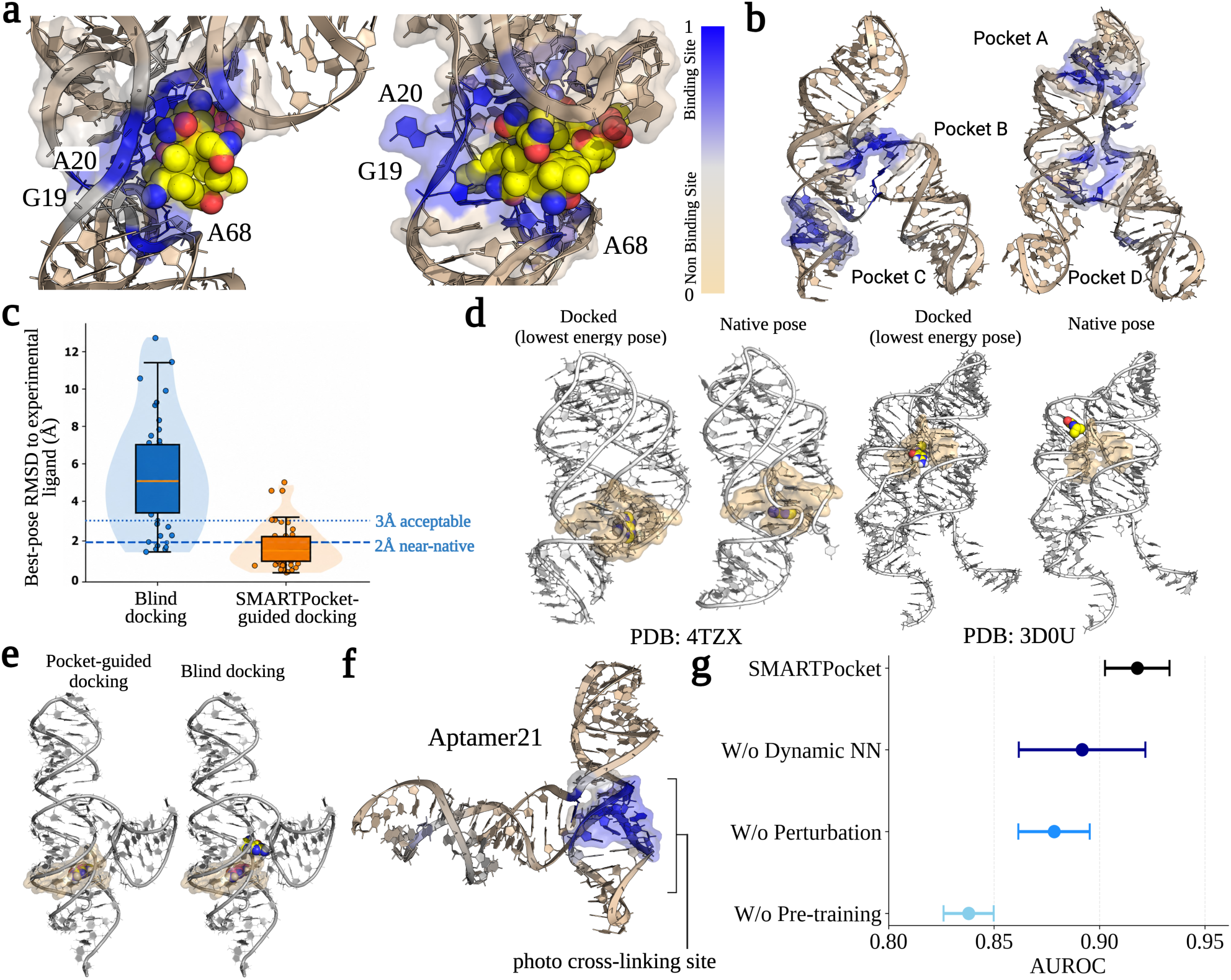
SMARTPocket identifies cryptic and experimentally validated RNA binding sites, guides molecular docking, generalizes to predicted RNA structures, and ablation studies. **(a)** Two orthogonal views of the cobalamin riboswitch with SMARTPocket-predicted binding residues (tan-to-blue gradient) accurately localizing the cryptic pocket. Key residues G19, A20, and A68 are labeled; the ligand is rendered in spheres. **(b)** SMARTPocket identified pockets A, B, C, and D on the SARS-CoV-2 RNA structure correspond to experimentally validated binding sites, illustrating the potential utility of SMARTPocket for therapeutic target identification. **(c)** Distribution of best-pose RMSD values comparing conventional blind docking and SMARTPocket-guided docking across benchmarked RNA–ligand complexes. SMARTPocket-guided docking substantially shifts docking solutions toward lower RMSD values, enriching for near-native ligand poses within the <2 Å and <3 Å accuracy thresholds. **(d)** Representative examples of binding pockets identified by the lowest energy docked pose (left) and the docked ligand adopted orientation to the native ligand. RNA surfaces are shown in transparent orange representation, while ligands are displayed as stick or sphere models colored by atom type. Across diverse RNA folds, SMARTPocket-guided docking consistently localizes ligands within experimentally observed binding pockets and recovers native binding geometries with substantially improved accuracy relative to unconstrained blind docking**. (e)** Comparison of SMARTPocket-guided and blind docking using the native ligand. Pocket-guided docking affords a near native pose with an RMSD less than 1 Å (left) while blind docking positions the native ligand in an altenative pocket close to the native pocket (right). **(f)** SMARTPocket identifies putative photo-crosslinking sites on the computationally predicted Aptamer21 structure, demonstrating applicability to modeled RNA structures lacking experimental coordinates. **(g)** SMARTPocket ablation study comparing the AUROC of the complete SMARTPocket model against variants lacking protein pretraining, coordinate perturbation augmentation, or dynamic nearest-neighbor graph construction, evaluated on the combined HARIBOSS, filtered Jiang, and time-dependent test sets. Ablated variants exhibit performance degradation, confirming the contribution of each component to overall model performance.

The SARS-CoV-2 FSE RNA, for which multiple ligand-binding pockets (A–D) have been experimentally mapped through Chem-CLIP and computational screening^43^, was next assessed. Using ten independently refined structural models of the FSE (PDB:6XRZ)^46^, SMARTPocket recovered Pocket A, the experimentally validated merafloxacin-binding site, in seven models, while identifying Pocket B in five models and Pocket C in three^43^. Pocket D was detected only once and with low confidence. These data demonstrate that SMARTPocket accurately captures multiple ligandable pockets directly from RNA structure, without requiring ligand information (Fig. 5b). Note that both the cobalamin riboswitch and SARS-CoV-2 FSE RNA exhibited low structural similarity to the training and validation sets (RMScore cutoff = 0.5), making these studies a blind testing scenario.

Together, these analyses demonstrate that SMARTPocket can identify ligandable RNA binding sites across diverse structural contexts, from cryptic pockets that emerge through conformational rearrangement to experimentally validated binding hotspots in viral regulatory RNAs. The ability to detect both pre-formed and cryptic pockets directly from RNA structure underscores SMARTPocket’s utility as a general framework for RNA-targeted small molecule discovery. Moreover, when paired with Chem-CLIP^47^, which uses covalent small molecule probes to capture ligand–RNA contacts and sequencing-based readouts to map those interactions at nucleotide resolution^48^, SMARTPocket can connect predicted ligandable pockets with experimentally observed sites of small-molecule occupancy. This combined computational and chemical mapping strategy can prioritize RNA binding pockets, validate pocket assignments, and guide ligand optimization without requiring ligand-bound RNA structures.

### SMARTPocket-guided docking improves RNA–small molecule pose prediction accuracy

We next sought to determine if SMARTPocket-guided docking improves pose prediction as compared to blind docking. In blind docking, the full RNA surface is searched without prior knowledge of the binding region^49^. This greatly expands the conformational search space and increases the likelihood of selecting energetically favorable but non-native poses^49^. Blind docking also requires a known ligand, typically the native ligand or a close analog, to infer potential binding regions. As a result, performance can be highly target-dependent, especially for large or conformationally heterogeneous RNAs^50^. SMARTPocket provides a complementary strategy by first identifying probable ligand-binding pockets directly from RNA structure, without requiring ligand information. Docking calculations can then be restricted to these predicted pockets. By narrowing the search space, we hypothesized that SMARTPocket-guided docking could increase the recovery of near-native poses, reduce high-RMSD poses, and improve docking consistency.

Across benchmark RNA–ligand complexes and structural motifs, including hairpins, bulges, internal loops, and junctions (Supplementary Table 1), SMARTPocket-guided docking using the AutoDock-GPU^51^ produced lower RMSD values relative to experimentally observed ligand poses than blind docking. The RMSD distributions were tighter and shifted toward near-native conformations, with a larger fraction of poses below the near-native threshold of <2 Å (Fig. 5c, e). In contrast, blind docking (lowest energy docked pose) often identified broad regions that overlapped geometrically with the native binding pocket, defined here as proximity within 10 Å of the native ligand center. However, approximate pocket localization did not reliably produce near-native ligand placement. Many blind-docking poses still deviated substantially from the experimentally observed binding mode, indicating that broad pocket detection alone is insufficient for accurate RNA–ligand pose recovery (Fig. 5d). In addition to improving pose accuracy, SMARTPocket-guided docking substantially increases computational efficiency. Preparation of blind docking calculations typically requires 30 seconds to several minutes per target, depending on RNA size, because the docking box must encompass the entire RNA structure. In contrast, SMARTPocket restricts docking to a predefined binding pocket, dramatically reducing the search space and the number of ligand configurations that must be evaluated. As a result, docking calculations are completed more rapidly, an advantage that becomes increasingly significant for large RNA targets and ultra-large virtual screening campaigns.

These results show that structure-based pocket prediction can improve RNA docking by focusing the search on ligandable regions before pose generation. By identifying RNA binding pockets directly from structure, SMARTPocket-guided docking reduces dependence on exhaustive surface searching, improves native pose recovery, and provides a more reliable workflow for RNA structure-based ligand discovery.

### SMARTPocket accurately identifies binding pockets on computationally predicted RNA 3D structures

Chemical mapping approaches including Selective 2’-Hydroxyl Acylation analyzed by Primer Extension (SHAPE)^52^, dimethyl sulfate (DMS) chemical probing^53^, Photoaffinity Evaluation of RNA Ligation-Sequencing (PEARL-seq)^54^, and Chem-CLIP-based mapping approaches^48^ have demonstrated that Aptamer 21^55^ contains a discrete small-molecule binding pocket for dihydropyrimidine ligands, despite the absence of an experimentally determined 3D structure. In these studies, ligand-dependent cross-linking, reverse-transcription stops, and sequencing-based mutational profiling localized small molecule engagement to a structured region within the three-way junction of the RNA^54^. Prominent signals were observed at C20 and A34, with additional reverse-transcription drop-off or mutation signals at C39 and G58, consistent with a compact binding site spanning the junction core and adjacent structural elements. Although these chemical mapping experiments identified the nucleotides involved in ligand recognition, the lack of high-resolution structural data prevented direct visualization of the pocket geometry in three dimensions.

To test whether SMARTPocket could identify ligandable pockets in computationally predicted RNA structures, an ensemble of de novo tertiary structures for Aptamer 21 using Fragment Assembly of RNA with Full-Atom Refinement (FARFAR2^22^) was generated, where FARFAR2 reconstructs stereochemically realistic RNA folds guided by experimentally constrained secondary structure^22^. Applying SMARTPocket to the FARFAR2-derived structural ensemble, with no experimentally derived restraints, revealed a single, high-probability ligandable pocket that localized to the region encompassing C20, A34, C39, and G58. This prediction closely matched the binding site defined by Chem-CLIP^48^,PEARL-seq^54^, SHAPE^52^, and DMS mapping^53^ (Fig. 5f). Notably, SMARTPocket required no ligand information, photo-cross-linking data, or experimental restraints beyond the predicted RNA structures themselves.

### SMARTPocket ablation study

Ablation studies were conducted to assess the contribution of each model component across the aggregated test samples from HARIBOSS, filtered Jiang, and the time-dependent test set (Fig. 5g). Removing protein data pretraining produced the largest performance drops, with AUROC declining from 0.918 ± 0.015 (mean ± s.d.) to 0.838 ± 0.012. These results highlight that large-scale protein pretraining equips the model with transferable geometric knowledge that effectively mitigates RNA–ligand data scarcity and enhances RNA binding site prediction.

In addition to cross-system transfer learning, a simple coordinate perturbation augmentation was utilized to augment the limited RNA-ligand training data. Removing this augmentation reduced model performance to an AUROC of 0.878 ± 0.017, underscoring its effectiveness. By generating four coordinate-perturbed variants per RNA, with slightly displaced atomic coordinates, the training set was expanded five-fold, providing an implicit regularization effect that improved model performance.

The dynamic message passing design, which exposes each atom to a variable number of nearest neighbors at each layer while progressively expanding neighborhood size with network depth, was also evaluated. Removing dynamic message passing resulted in a performance decrease, with AUROC dropping to 0.893 ± 0.030. This result suggests that stochastic neighborhood sampling helps reduce overfitting by providing an additional form of model regularization.

## DISCUSSION

In this study, we introduce SMARTPocket, a geometric deep learning model that advances RNA–small molecule binding pocket prediction by directly leveraging atomic-level information. Built upon a geometric transformer^18^ architecture, SMARTPocket learns 3D geometric patterns of molecular recognition without relying on conventional handcrafted structural features. The design ensures rotational equivariance and translational invariance in binding pocket prediction and scales naturally to multi-chain RNA assemblies, addressing key limitations of most existing approaches.

A second key innovation is SMARTPocket’s cross-system transfer learning strategy. Because experimentally solved RNA–small molecule complexes remain limited, SMARTPocket uses the larger body of protein structural data to learn general atomic interaction patterns. These features are then adapted to RNA-specific structural patterns during fine-tuning. Ablation studies show that removing the pretraining step substantially reduces performance, indicating that cross-domain transfer is not only feasible but important for accurate RNA–small molecule pocket prediction.

SMARTPocket fits into a growing number of computational approaches for RNA-targeted small-molecule discovery. Motif- and sequence-based platforms such as Inforna^56^ and SMARTBind^15^ are powerful because they can prioritize RNA–ligand interactions without requiring experimentally determined three-dimensional structures. These approaches are especially useful when only RNA sequence or secondary structure information is available. However, they do not directly define the three-dimensional shape, enclosure, or atomic composition of a ligand-binding pocket, which are critical for structure-based ligand design and docking. In contrast, structure-based tools such as fpocketR^16^, DRLiPS^42^, SHAMAN^17^, and docking-based workflows explicitly evaluate RNA three-dimensional structure. These methods provide complementary capabilities: fpocketR identifies geometric cavities, DRLiPS estimates RNA pocket druggability from structure-derived features, SHAMAN evaluates small-molecule binding sites across RNA conformational ensembles, and blind docking can identify ligand poses when a candidate ligand is already available. SMARTPocket is complementary to these approaches because it predicts ligandable nucleotides directly from RNA structure without requiring a known ligand, handcrafted pocket descriptors, or exhaustive docking.

Across benchmark datasets, SMARTPocket consistently exhibits strong predictive performance. Notably, SMARTPocket reliably identifies ligand-binding pockets from apo structures when conformational differences are limited, though performance decreases in cases where ligand binding induces major structural reorganization. Beyond canonical pockets, SMARTPocket also identifies cryptic binding sites, as demonstrated in cobalamin riboswitch, where the model detects ligandability before full pocket opening^21^. SMARTPocket also recapitulates experimentally validated binding regions in the SARS-CoV-2 frameshifting element^43^ and recovers chemically mapped pockets in computational models predicted by FARFAR2^22^ of Aptamer 21. These results show that SMARTPocket can extend pocket prediction to RNAs lacking ligand-bound structures, broadening the scope of RNA targets that can be evaluated for small-molecule recognition.

SMARTPocket is also complementary to experimental RNA-target mapping methods such as Chem-CLIP^47^. Chem-CLIP uses covalent small molecule probes to capture direct ligand–RNA interactions and map binding sites through reverse-transcription stops, sequencing, or mutational readouts^48^. These data can define the nucleotides contacted by a ligand, but they do not by themselves provide a three-dimensional model of the pocket. SMARTPocket fills this gap by projecting chemically mapped binding information onto predicted or experimental RNA structures, allowing nucleotide-level Chem-CLIP signals to be interpreted as three-dimensional ligandable pockets. This combined strategy can help prioritize RNA binding sites, guide analog design, and support structure-informed optimization even when RNA structures are unavailable.

This concept builds on earlier RNA-targeting strategies from our laboratory, including Inforna^56^ and sequence- or motif-based design approaches^15^, which linked RNA sequence and secondary structure motifs to small molecule recognition, as well as sequence-informed deep learning models such as SMARTBind^15^. SMARTPocket extends this framework from two-dimensional motif recognition to three-dimensional pocket prediction. In this way, it connects sequence-informed RNA ligand discovery, experimental occupancy mapping by Chem-CLIP, and atomic-level structural modeling into a unified strategy for identifying and optimizing RNA-targeted small molecules.

Despite these advances, SMARTPocket has limitations. First, processing very large RNA assemblies, such as ribosomal subunits or spliceosomes, remains computationally demanding because full-atom attention operations scale poorly with system size. Future versions may benefit from hierarchical atom-to-residue modeling and more efficient attention mechanisms. Second, SMARTPocket currently operates on static structural snapshots and does not explicitly model RNA dynamics, even though conformational plasticity is central to RNA ligand recognition^40^. Incorporating molecular dynamics-derived ensembles^57^ or experimentally constrained structural ensembles may further improve prediction of transient and cryptic pockets.

In summary, SMARTPocket provides a robust and generalizable framework for identifying ligandable RNA pockets directly from structure. By detecting canonical, apo-accessible, chemically mapped, and cryptic pockets on both experimental and computationally predicted structures, SMARTPocket addresses a central challenge in RNA-targeted drug discovery. When integrated with Chem-CLIP and sequence-informed platforms such as Inforna, SMARTPocket provides an experimentally grounded, structure-guided path for discovering and optimizing RNA-targeted small molecules. Ultimately, however, compounds emerging from these and other computational approaches will require rigorous medicinal chemistry optimization to convert initial binders into useful chemical probes and therapeutic leads — an iterative process that tools like SMARTPocket are designed to inform and accelerate.

## ACKNOWLEDGEMENT

This work was supported in part by the University of Florida (UF Startup Fund, UF Health Cancer Institute Pilot Grant # UFS-2023-08, and UF-NVIDIA Artificial Intelligence and Complex Computational Research Award to Y. L.), National Institutes of Health (R21EB037868 to Y. L. and R01 CA249180 to M.D.D.), the Florida Department of Health (Florida Cancer Innovation Fund grant #MOABO to J. L. C.), the Nicholas Bodor Professorship Fund (to C. L.), and the Muscular Dystrophy Association (Development grant ID #963835 to A. T.). We thank Reza Esmaeeli (University of Florida) for technical advice and discussions. Figures were created in Biorender https://BioRender.com.

## Methods

### Dataset curation

#### Single-chain RNA benchmark datasets

The TR60 training set and four associated benchmark test sets (TE18, RB9, TL12, and JL10), which are widely used in RNA-small molecule binding site prediction studies, consist solely of single-chain RNA data. The TR60 dataset was curated by RNASite^9^ study, containing RNA-small molecule complexes where the ligands are small molecules with molecular weights ≤ 1000 Daltons. These ligands include metal ions, natural metabolites, and drug-like molecules, but exclude water molecules and crystallization additives. The RNA structures were filtered for redundancy using TM-scoreRNA^33, 59^ for pairwise structure similarity. Specifically, when the TM-scoreRNA exceeded 0.3, the structure with the highest number of binding-site nucleotides was retained. The resulting 78 RNAs were clustered at 30%-similarity using CD-HIT-EST^60^ and BLASTclust^61^, yielding a training set of 60 RNAs (TR60) and a test set of 18 RNAs (TE18). The training set was subsequently split 9:1 into training and validation subsets.

RB9 is a benchmark dataset derived from the RBind^7^ study, originally comprised 19 RNA–small molecule complexes. After removing entries exhibiting overlap with the TR60 training set, a final set of 9 RNA–ligand complexes were retained^9^. JL10 and TL12 are two additional benchmark datasets introduced in the ZHmolReSTasite^10^ study to further expand the scope of available evaluation datasets. JL10 contains 10 structurally complex RNAs containing junction loops, selected from high-resolution PDB structures (resolution < 4Å) released after January 2021, filtered to exclude protein/DNA interactions, structures with > 2 junction loops or with < 10% binding nucleotides, and redundant samples with redundancy removed at 95% sequence similarity. TL12 contains 12 low structure complexity RNA chains interacting with ligands, collected after January 2021, specifically excluding junction loop-containing structures to contrast with JL10’s complexity.

#### Broader curated RNA datasets

RNA–ligand complexes frequently involve multiple chains that together coordinate a single ligand, forming binding sites that depend on contributions from each chain. To capture the full complexity of ligand-binding environments and establish robust benchmarking of binding site prediction methods, three datasets encompassing both single-chain and multi-chain RNA structures were assembled, ensuring accurate and complete representation of binding pockets.

#### HARIBOSS dataset

A major benchmark dataset was derived from the HARIBOSS database, which, at the time of this study, comprises 868 RNA–ligand complexes and 1,471 annotated binding pockets. The following filtration steps were applied: (1) exclusion of ribosomal RNA and (2) removal of complexes where the small molecule did not form a binding pocket (e.g., end-stacking interactions) or in which ligand was a non-interested drug-like small molecule. This procedure yielded 447 RNA structures. For structures with multiple models, the first one was retained.

For data splitting, pairwise structural similarity was first assessed using RMScore, and hierarchical agglomerative clustering was performed with a cutoff of 0.5, yielding 25 clusters. Clusters were assigned to splits such that no test sample shared an RMScore above 0.5 with any training or validation structure, resulting in 357 training, 32 validation, and 58 test RNA structures. To minimize data leakage, an additional pocket-level similarity check was performed, defining each binding pocket as the set of residues within 4 Å of the ligand and applying the same RMScore cutoff of 0.5 between test and training/validation pockets, reducing the final test set to 44 structures.

#### Filtered Jiang test set

The original Jiang dataset**^Error! Reference source not found.^**^36^ comprises 800 NA-ligand structures. Subsequent filtering removed non-drug-like (hydrogen bond donors < 4, molecular weight > 110, hydrogen bond acceptors < 6, rotatable bonds < 4, chiral centers > 0, AlopP <4, polar surface area < 110) and complexes with crystal contact artifacts, yielding a set of 255 structures^36, 62^. In this study, additional filtering steps were applied to ensure dataset quality and non-redundancy. First, DNA-ligand complexes, structures overlapping with the HARIBOSS dataset, and complexes containing buffer molecules as ligands (eg. MES, EPE) were excluded. Next, structures with RNA and pocket structure similarity above RMScore 0.5 relative to the HARIBOSS training and validation sets were removed to prevent data leakage. Finally, structurally redundant RNAs within the test set (pairwise RMScore >0.8) were removed, resulting in a final curated dataset of 6 representative RNA-small molecule complexes.

#### Time-dependent test set

This dataset comprises RNA-small molecule complexes from the PDB released after September 30, 2021, ensuring evaluation of structures not present in the AlphaFold3 training dataset. A similar filtering process was applied to remove ribosomal RNA structures, complexes present in previous benchmarks, and complexes in which small molecule did not form a binding pocket. Samples exhibiting RNA and pocket structure similarity above RMScore 0.5 relative to the HARIBOSS training and validation sets were excluded. A threshold (RMScore >0.8) was applied to remove redundant structures within the test set itself. This procedure yielded a final test set of 10 structurally diverse RNA-small molecule complexes.

For all above compiled datasets, water molecules and hydrogen atoms were removed from the RNA-small molecule complexes. The binding site nucleotides are defined as nucleotides containing at least one atom within 4 Å of any atom in the bound ligand.

### SMARTPocket Model

#### Features

The RNA structure input is represented as an atomic point cloud, with each atom described by its three-dimensional coordinates and element type. Structural geometry is encoded using pairwise distances and normalized displacement vectors, which ensure translation invariance. Specifically, for each atom *i* with spatial coordinates *x*_j_ ∈ ℝ^3^ and its *n* nearest neighbors *j* ∈ *N* (*i*) with coordinates *x*_j_ ∈ ℝ^3^, the pairwise distances 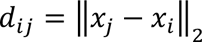 and normalized displacement vectors 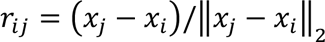 are computed to characterize the local geometric environment of central atom *i*. An *n*-nearest-neighbor graph is constructed for each atom, where each node *i* maintains two states: a scalar state *q*_i_ ∈ ℝ^5^ and a vector state 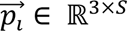, where *S* denotes the state dimension. The node features are constructed as 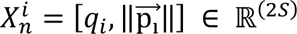, and the edge features are defined as

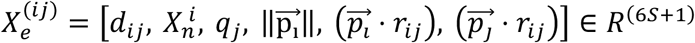

where 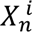 represents the central node *i*’s features, *q_i_* and ∥ *p_j_*∥ capture the neighbor *j*’s scalar and vector features respectively, and the projection terms *p*_i_ ⋅ *r*_ij_ and *p*_j_ ⋅ *r*_ij_ measure the alignment of center and neighbor vector states with the edge direction, respectively, and square brackets [·] denote the concatenation operation.

#### Model architecture

Our model builds on PeSTo’s geometric transformer architecture and extends to the task of RNA binding pocket prediction. This design was motivated by PeSTo’s strong ability to predict protein binding interfaces at atomic resolution, providing a robust model foundation and transferable geometric and interaction priors for RNA structure modeling. Each geometric transformer consists of two coupled attention-based modules with key-query-value components for updating the scalar and vector states of each node. Queries representations are derived from the node features *X*_n_, which are passed through a multilayer perceptron (MLP) to produce scalar queries 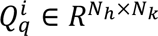 and vector queries 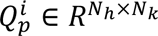, where *N_n_* denote the number of attention heads and *N* specifies key-query dimension. Keys and values are derived from the edge features *X*_e_ via separate MLPs, each comprising scalar and vector components: scalar keys 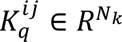 and vector keys 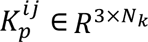, as well as scalar values 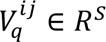and vector values 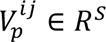.

The resulting attention-weighted messages are subsequently aggregated to update the scalar and vector node states, as formalized in the following equations

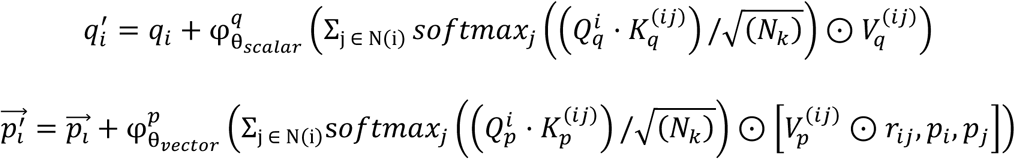

where 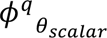 and 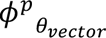 are MLPs.

With increasing depth of the geometric transformer layers, the number of nearest neighbors *n* was progressively expanded following a pre-defined configuration, allowing each central node to incorporate broader structural context through an enlarged neighborhood *N*(*i*). Meanwhile, to enhance model generalizability, a random integer perturbation sampled from a normal distribution with standard deviation of 2 was added to the pre-defined neighbor count at each layer, enabling a dynamic nearest-neighbor graph construction and message-passing process during RNA data fine-tuning.

To aggregate atomic representations into nucleotide-level features, a self-attention–based pooling mechanism was applied over all atoms within each nucleotide to compute the corresponding nucleotide-level representation.

Finally, a decoder MLP classifies each nucleotide as part of the binding interface, producing probability scores between 0 and 1 that indicate the likelihood of nucleotide involvement in ligand binding.

#### Model training

To address the limited availability of RNA structure data for model training, a simple data augmentation strategy was employed. The training sets were augmented by including both the original RNA structures and coordinate-perturbed variants. In the perturbed structures, atomic coordinates along the x, y, and z axes were independently and randomly modified by 5% of their respective standard deviations. This procedure increases the sizes of the two RNA-ligand training sets, which originally contained only 54 and 357 structures, by approximately fivefold, thereby helping to mitigate model overfitting.

The model was trained using the AdamW^63^ optimizer with an initial learning rate of 1 × 10⁻⁴, tuned empirically to balance convergence speed and training stability. A stepwise learning rate decay (StepLR scheduler, step size = 7, γ = 0.5) was applied. The loss function was binary cross-entropy, augmented with class re-weighting to address class imbalance. Meanwhile, label smoothing^64^ with a factor of 0.15 was applied to reduce the prediction overconfidence and enhance generalization. Early stopping was applied based on minimum validation loss, with a patience of 5 epochs, terminating training if no improvement was observed within that window. All training was performed using PyTorch with CUDA acceleration on a single NVIDIA A100 GPU with 80 GB Memory.

#### Baseline Methods

In this study, SMARTPocket was benchmarked against 13 recently published and classical RNA-focused binding site predictors, as well as the general-purpose molecular complex structure prediction model. Based on their methodologies, these methods can be roughly categorized into four classes. The first category comprises classical geometry and network topology-based methods, including Rsite^5^, Rsite2^6^, RBind^7^, and fpocketR^16^, which identify binding sites through distance measures, graph centrality metrics, or cavity detection algorithms. Rsite predicts RNA functional sites by computing Euclidean distances between each nucleotide and all other nucleotides in the RNA tertiary structure, identifying nucleotides at the extreme points of the resulting distance curve as candidate binding sites. Rsite2 extends this distance-based approach to RNA secondary structure, replacing tertiary structure derived Euclidean distances with Hamming distances computed from predicted secondary structure coordinates, thereby eliminating the requirement for experimentally determined three-dimensional structures. RBind converts RNA tertiary structures into nucleotide interaction networks and applies degree and closeness centrality measures to identify ligand and protein binding nucleotides. fPocketR is a recently published RNA-optimized pocket detection method adapted from geometry-based fpocket framework for identifying ligandable pockets in RNA tertiary structures.

The second category consists of feature-engineering-based machine learning or deep learning methods, including RNASite^9^, RNET^8^, RLBind^11^, and ZHMolReSTasite^10^, which extract handcrafted structural, network, or surface topography descriptors and feed them into machine learning classifiers for RNA binding site prediction. RNAsite uses a random forest classifier that combines sequence profiles derived from multiple sequence alignments with structure-based descriptors to predict RNA binding sites. RNet builds on structural network features and uses an ensemble of machine learning classifiers, including random forest, LightGBM, and XGBoost, for RNA binding sites and dynamical binding behavior prediction. RLBind employs a dual-channel convolutional neural network that integrates global and local features derived from full-length and local neighbor RNA sequence and structure properties to predict RNA–ligand binding sites. ZHMolReSTasite introduces RNA surface topography by generating topographic images from solvent-excluded surfaces of RNA tertiary structures and integrating sequence, structural, geometric and environmental features. A convolutional neural network is then used to predict binding nucleotides.

The third category encompasses recent data-driven deep learning-based methods, including RNABind^12^, GATRSite^13^, SMARTBind^15^, and GerNA-Bind^14^, which bypass manual feature engineering and instead learn complex sequence or structure relationships underlying ligand recognition directly from data. RNABind and GATRSite combine pretrained RNA language model sequence-based embeddings with graph-based structural representations to predict binding sites. GerNA-Bind integrates multi-state representations of RNA and ligands across sequence, secondary structure and three-dimensional conformations using geometric deep learning to jointly predict binding specificity and binding sites. SMARTBind is a structure-agnostic RNA-targeted ligand discovery framework that leverages RNA language models, contrastive learning and ligand-specific decoy enhancement to predict RNA–ligand binding scores and binding sites directly from primary RNA sequences.

The fourth category consists of general-purpose biomolecular structure prediction models, represented here by AlphaFold3^20^, which predicts RNA–ligand binding sites indirectly by co-folding RNA sequences with ligand SMILES and extracting binding nucleotides from the resulting predicted complex structure.

#### Evaluation metrics

Binding site prediction performance was evaluated using six metrics, including threshold-independent measures, such as area under the receiver operating characteristic curve (AUROC) and area under the precision-recall curve (AUPRC), as well as threshold-dependent metrics, such as Matthews correlation coefficient (MCC), F1 score, precision and recall.

Precision measures the proportion of correctly predicted binding nucleotides among all positive predictions. Recall, also known as sensitivity, quantifies the fraction of true binding nucleotides successfully identified by the model. F1 score is the harmonic mean of precision and recall. MCC is a balanced measure that considers true and false positives and negatives, providing a single value that summarizes the confusion matrix. Unlike the decision threshold-dependent metrics, AUROC measures how accurate the model is at ranking binding nucleotides above unbinding ones at all cutoffs. AUPRC accesses how well the model balances precision and recall over varying thresholds. The formulas for the metrics are given by:

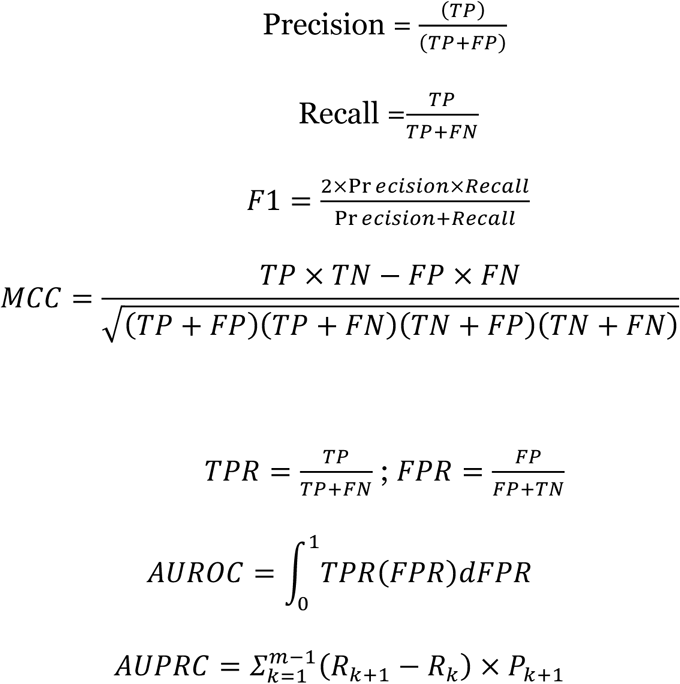

where TP denotes true positives (correctly predicted binding nucleotides), TN true negatives (correctly predicted non-binding nucleotides), FP false positives (non-binding nucleotides incorrectly predicted as binding), and FN false negatives (binding nucleotides incorrectly predicted as non-binding). *R*_k_ and *P*_k_ are the recall and precision and the k-th threshold.

### Data and Code availability

RCSB PDB (https://www.rcsb.org/), HARIBOSS (https://hariboss.pasteur.cloud), and single-chain RNA benchmark (http://zhaoserver.com.cn/ZHmol/ZHmolReSTasite/ZHmolReSTasite.html) datasets are publicly available. Source code for SMARTPocket is publicly available via GitHub at https://github.com/AIDD-LiLab/SMARTPocket. The benchmark models are available from their official GitHub repositories and web pages: RNASite (https://yanglab.qd.sdu.edu.cn/RNAsite/), RLBind (https://github.com/KailiWang1/RLBind/tree/ab6d1cc1324f05da58bd246cc660c7a64b29ba37), fpocketR (https://github.com/Weeks-UNC/fpocketR/tree/f397fe596f850e93bd1449e8705b05e0bcb84063), GerNA-Bind (https://github.com/GENTEL-lab/GerNA-Bind/tree/33e93139432cdc194368b2254d628750b714450f), RNABind (https://github.com/jaminzzz/RNABind/tree/7936279ff92a929a2de382ac902b9f1021c6c46b), SMARTBind (https://github.com/AIDD-LiLab/SMARTBind/tree/350fbfbbf60a89401499b8ee5119914c74b44ff0), AlphaFold3 (https://github.com/google-deepmind/alphafold3/tree/bc73f5652b40599bc40fd18215d8190f20b15d8f).

## SUPPLEMENTARY INFORMATION

Supplementary Information includes Supplementary Notes detailing dataset information and molecular docking implementation, Supplementary Figures 1-8, Supplementary Tables 1 –9, and additional Experimental Methods.

