## Supplementary Information for "Geometric Deep Learning Reveals Ligandable and Cryptic RNA Binding Small Molecule Pockets (SMARTPocket)"

### SUPPLEMENTARY FIGURES & TABLES

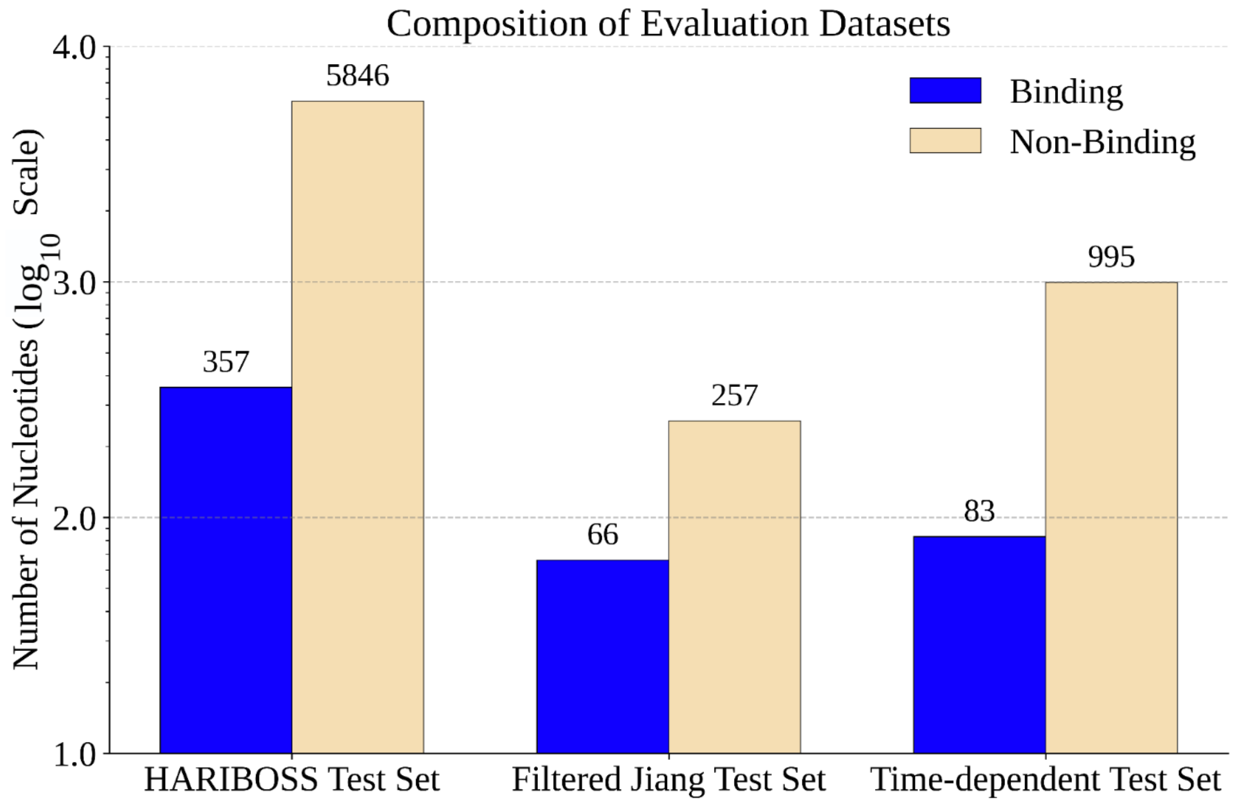


**Supplementary Fig. 1 | Distribution of binding and non-binding nucleotides across the three broader curated test sets.**

Bar plot showing the number of binding (blue) and non-binding (beige) RNA nucleotides in three broader curated test datasets: HARIBOSS[1], filtered Jiang[2], and time-dependent test sets. Counts are shown on a $\log_{10}$ scale. Binding nucleotides are substantially outnumbered by non-binding across all datasets.

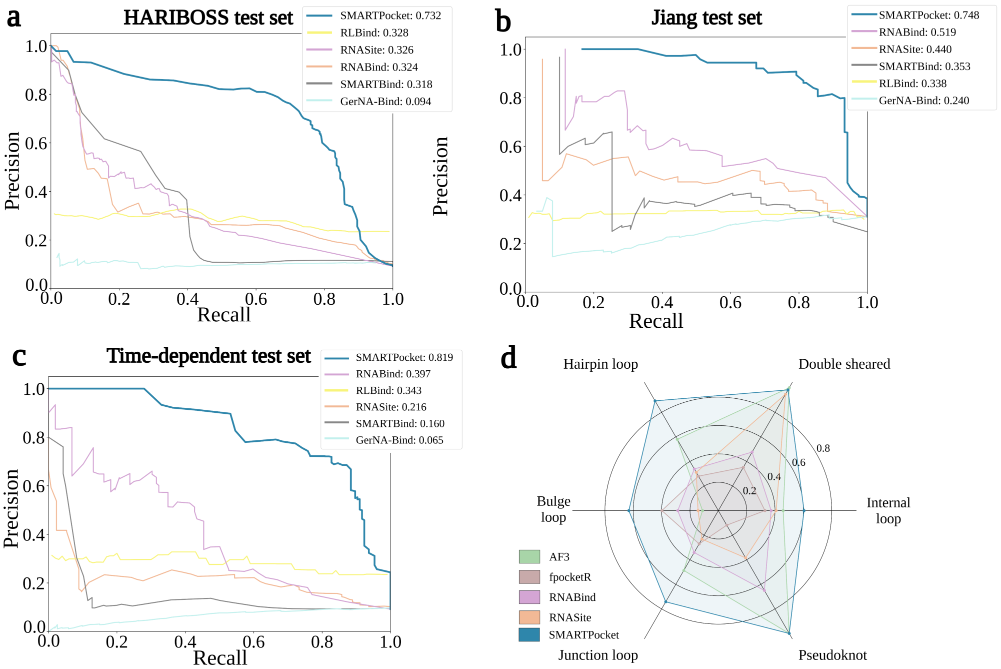


**Supplementary Fig. 2 | Precision-recall curves and motif comparison for SMARTPocket and baseline models on three broader curated test sets.**

**(a)** Precision-recall (PR) curves on the HARIBOSS[1] test set wherein SMARTPocket achieves an AUPRC of 0.732 ± 0.033 and compared with five baseline methods. **(b)** PR curves on the filtered Jiang[2] set, where SMARTPocket achieves an AUPRC of 0.748 ± 0.041. **(c)** PR curves on the time-dependent test set, where SMARTPocket achieves an AUPRC of 0.819 ± 0.052. In all 3 sets, SMARTPocket demonstrates performance superiority maintaining precision values exceeding 0.8 across a broad range of recall thresholds, indicating robust discrimination of true binding sites with minimal false positive rates. In contrast, competing methods exhibit precision decay at lower recall values, suggesting reduced specificity. **(d)** AUPRC values compared across structural motif classes annotated using RNA3DHub[3], evaluated on the combined HARIBOSS, filtered Jiang, and time-dependent test sets.

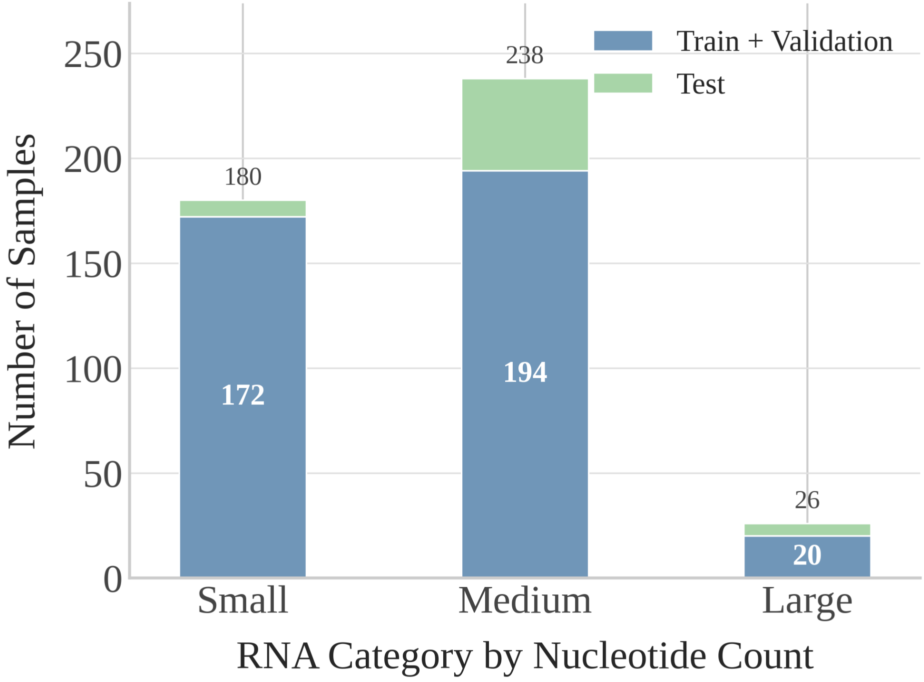


**Supplementary Fig. 3 | Distribution of RNA samples by nucleotide count in the HARIBOSS benchmark dataset.**

Bar plots showing the distribution of RNA samples by total nucleotide counts in the HARIBOSS benchmark dataset. The numbers of nucleotides are categorized as short (<50 nt; mean=34.6 ± 10.6 nt; n = 182), medium (50–200 nt; mean=89.1 ± 31.3 nt; n = 239), and long (>200 nt; mean=409.7 ± 60.8 nt; n = 26). Blue indicates the sample distribution for both training and validation sets, and green denotes the samples in the test set.

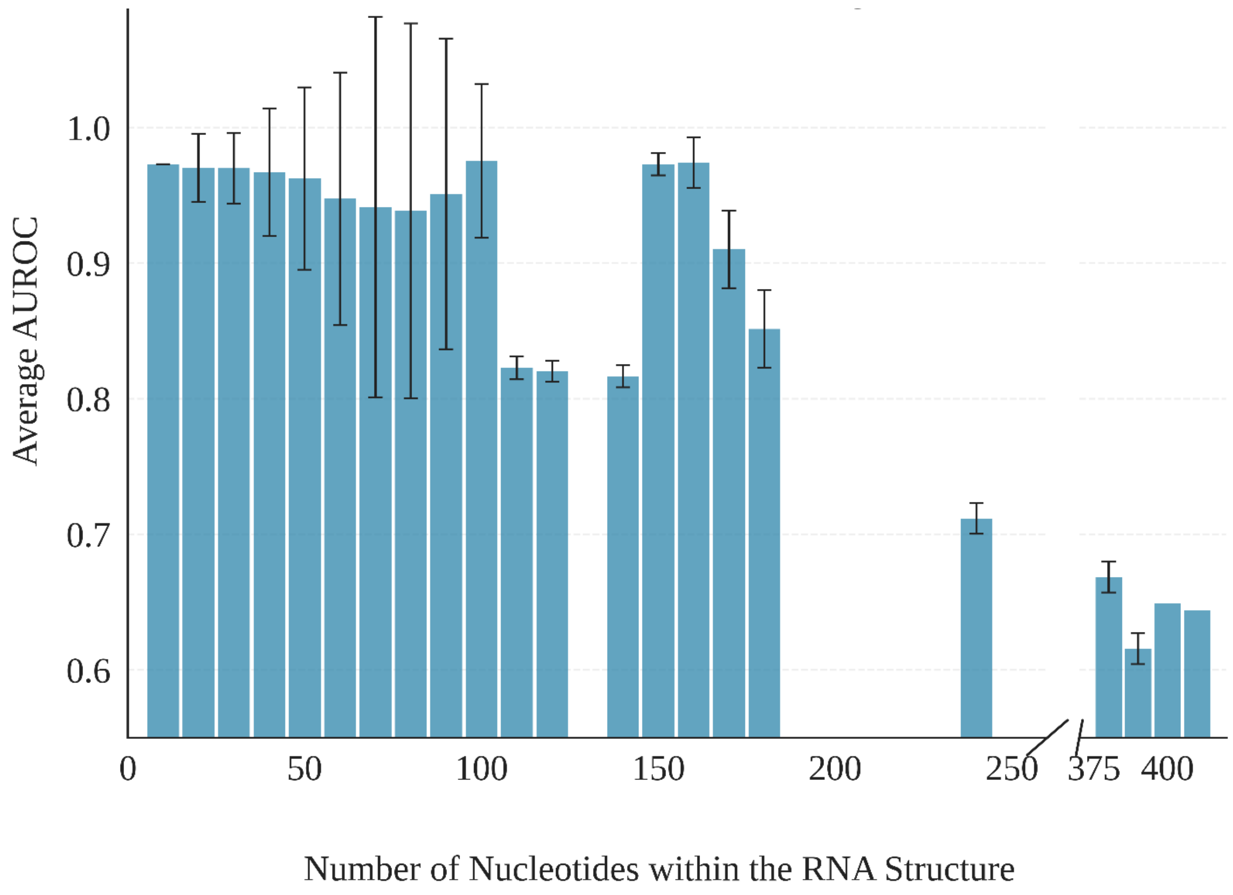


**Supplementary Fig. 4 | AUROC performance of SMARTPocket across RNA test cases grouped by nucleotide count.**

Histograms showing the average AUROC performance of SMARTPocket across RNAs (n=60), aggregated from three test datasets: the HARIBOSS[1], filtered Jiang[2], and the time-dependent test sets. RNA structures are grouped by nucleotide count using a bin size of 10. Error bars indicate the standard deviation across samples. SMARTPocket demonstrates consistently strong performance (AUROC > 0.8) on RNA structures with fewer than ~200 nucleotides. Performance declines for larger structures (≥375 nucleotides); however, the limited number of test cases between 200 and 375 nucleotides precludes precise characterization of the model performance within this range.

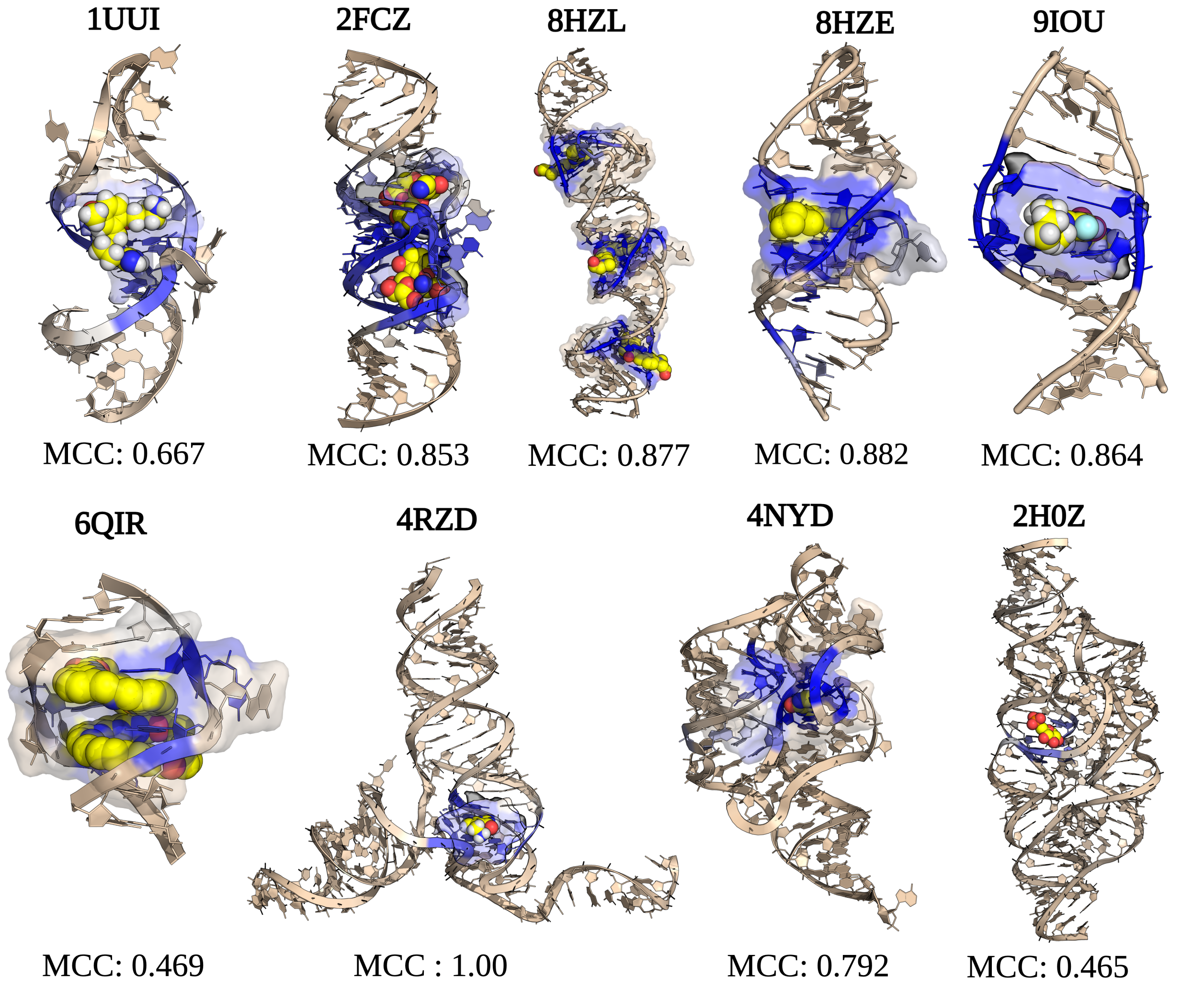


**Supplementary Fig. 5 | Visualizations of SMARTPocket predictions for representative RNA test cases.**

SMARTPocket-predicted binding sites for representative structures from test sets. Ligands are rendered as spheres, semi-transparent surfaces delineate ground-truth binding sites. Predicted binding confidence is encoded by a tan-to-blue gradient (tan, non-binding; blue, binding). MCC values are shown below each structure.

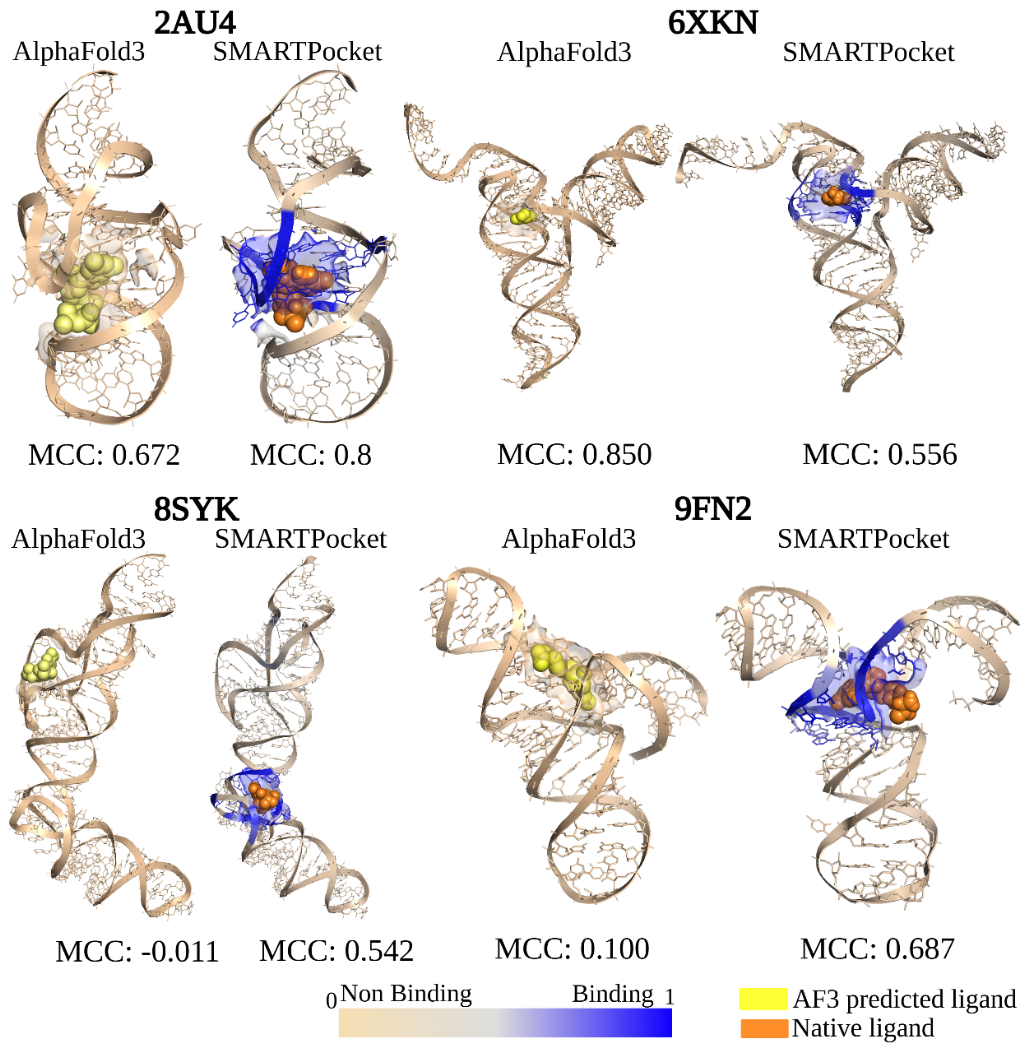


**Supplementary Fig. 6 | Visualization comparison of SMARTPocket and AlphaFold3 predictions for representative RNA test cases.**

Four representative case studies comparing binding site predictions from SMARTPocket and AlphaFold3[4]. AlphaFold3 performs strongly on the HARIBOSS test cases (2AU4 and 6XKN), where 2AU4 structure was released before model training data cutoff of September 30, 2021, suggesting data overlap; in contrast, performance drops on the time-dependent examples (8SYK and 9FN2), which fall outside this window. By contrast, SMARTPocket maintains consistently strong performance across both evaluation settings, demonstrating robust generalization to recently released, unseen RNA structures.

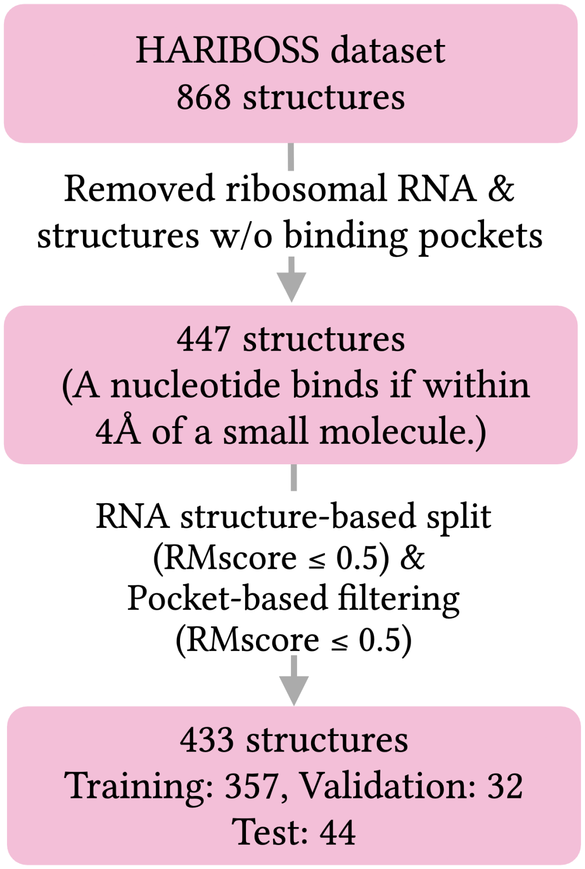


**Supplementary Fig. 7 | Overview of the dataset filtering workflow for the HARIBOSS benchmark dataset.**

Starting from 868 experimentally resolved RNA–ligand complexes curated in the HARIBOSS study[1], ribosomal RNAs, structures lacking defined binding pockets, and complexes with non-interested small molecules, such as water molecules and ions, were removed, resulting in 477 complexes. To ensure robust generalization assessment, the dataset was first split such that no test sample shared global RNA structural similarity with training or validation samples above an RMScore[5] cutoff of 0.5. An additional pocket-level similarity filter using the same cutoff was then applied to remove test samples with pockets similar to those in the training and validation sets. This procedure yielded 357 training, 32 validation and 44 test samples, comprising 433 structures in total.

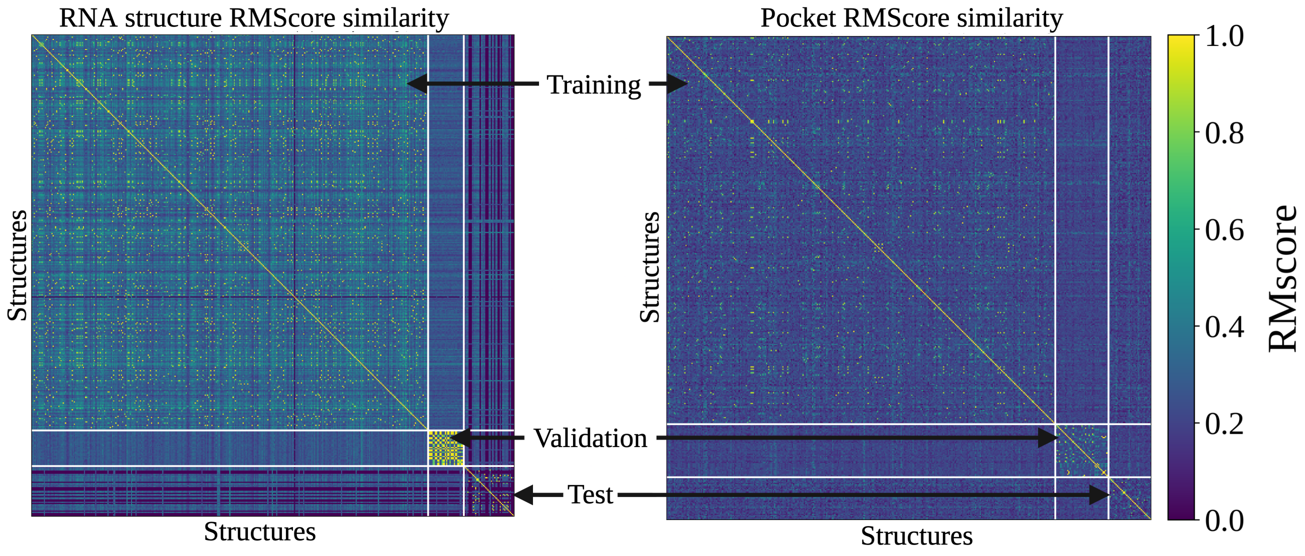


**Supplementary Fig. 8 | Pairwise RMScore similarity heatmaps for global RNA structure and local binding pocket across HARIBOSS data splits.**

Pairwise RMScore similarity was computed across all RNA structures in the HARIBOSS benchmark dataset. Because RMScore is asymmetric, values for each pair, (A, B) and (B, A), were averaged to obtain a symmetric similarity measure. Hierarchical agglomerative clustering yielded 25 clusters, which were used to partition structures into training, validation, and test sets. Left, pairwise RMScore similarity between global RNA structures. Right, pairwise RMScore similarity between local binding pockets, defined by nucleotides with any atom within 4 Å of a ligand atom. Colors indicate pairwise structural similarity, ranging from dark blue (RMScore = 0.0) to yellow (RMScore = 1.0). The consistently low RMScore values (dark blue) in the off-diagonal blocks between training, validation, and test sets confirm no structural overlap at either the global RNA structure or local pocket level, ensuring rigorous data partitioning.

**Supplementary Table 1 | List of PDB identifiers for single-chain RNA benchmarking datasets.**

| | | | | | | | | |
|---|---|---|---|---|---|---|---|---|
| TR60[6] | 1UUD | 1AJU | 4NYB | 3C44 | 1LVJ | 2L8H | 3TZR | 5FJ1 |
| | 1ARJ | 1Y90 | 3FU2 | 4PCJ | 1NBK | 2LWK | 3VRS | 5J02 |
| | 1BYJ | 1YLS | 3GX3 | 4QJH | 1NTB | 2MXS | 4FRG | 5O69 |
| | 1EHT | 1YRJ | 3MEI | 4QLM | 1QD3 | 2NOK | 4K31 | 5U3G |
| | 1EI2 | 2FCY | 3OXE | 4RGE | 1RAW | 2QUW | 4LVX | 5VCI |
| | 1F1T | 2G5K | 3Q3Z | 4XW7 | 1TOB | 2YIE | 4MEG | 6FZ0 |
| | 1FYP | 2KGP | 3Q50 | 5D5L | 1UTS | 3BNQ | 4MGM | 6HAG |
| | 1HR2 | 2KTZ | 3SKT | 5DH8 | | | | |
| TE18[6] | 1DDY | 2JUK | 4F8U | 379D | 1FMN | 2PWT | 430D | 4PQV |
| | 1F1T | 4YAZ | 5BJO | 6EZ0 | 1NEM | 5V3F | 364D | 2MIS |
| | 1Q8N | 2TOB | | | | | | |
| RB9[7] | 1NEM | 2GDI | 1Q8N | 3D2X | 1PBR | 2TOB | 1UUI | 3GX2 |
| | 1Y26 | | | | | | | |
| JL10[8] | 6WTR | 7KVU | 7UZ0 | 7UQ6 | 6WZS | 7TZR | 7EQJ | 7KGA |
| | 7EAF | 9BUN | | | | | | |
| TL12[8] | 7D12 | 7RWR | 7EOG | 8F0N | 7ELQ | 7TD7 | 7L0Z | 8FZA |
| | 7OA3 | 8HB1 | 7REX | | | | | |

**Supplementary Table 2 | List of PDB identifiers for HARIBOSS benchmarking dataset.**

**Training Set**

| | | | | | | | | | | | |
|---|---|---|---|---|---|---|---|---|---|---|---|
| 6DME | 1EHT | 3FO4 | 2BE0 | 8HBA | 6E82 | 3FO6 | 5FK2 | 6TFG | 5OB3 | 7D7Y | 1TN1 |
| 1AKX | 2M4Q | 6UET | 2A04 | 5Z71 | 1J7T | 6CB3 | 7FHI | 7RWR | 1KOC | 8U5K | 2HOJ |
| 2YGH | 6CK4 | 6HAG | 7EOP | 7EDT | 3SKR | 6CC1 | 2KX8 | 6E8T | 3IQR | 2GIS | 4Y1I |
| 3BNQ | 2F4S | 7D7V | 8G8Z | 3SLQ | 6E1S | 2L1V | 2N0J | 8D2B | 6LAX | 6DB8 | 3NPN |
| 7TDA | 6DLR | 4YAZ | 1F1T | 3SKW | 6VA4 | 7D7Z | 6YMJ | 3NPQ | 8D2A | 7D7X | 8R63 |
| 3FU2 | 6YMM | 7KVU | 5FKG | 4Q9R | 6T3R | 8R62 | 7U0Y | 3GX2 | 3IWN | 5FJC | 3V7E |
| 3SKI | 3RKF | 6P2H | 1FYP | 2HOP | 2ESJ | 5FKD | 7EDM | 1UUI | 5XI1 | 4TS2 | 5Z1I |
| 2ET3 | 6DLQ | 8D5O | 7FJ0 | 7WII | 2OE8 | 6TF0 | 3K0J | 6UC7 | 8U5T | 1NTA | 5ZEI |
| 8EYU | 7SZU | 4LX6 | 2KTZ | 5FKF | 4FAQ | 6LAZ | 1TN2 | 8U5Z | 2LWK | 4E8Q | 1FMN |
| 3TD1 | 2O3V | 2KU0 | 5FK6 | 1ARJ | 2HOO | 7D12 | 3SKZ | 7OAV | 2G5K | 3MUR | 3MUM |
| 8EYV | 6DLT | 3GCA | 7KVT | 6UC8 | 3GX5 | 4GPW | 7EON | 4KQY | 5FK5 | 6C64 | 6HC5 |
| 4YB0 | 3SKT | 3E5F | 7TZU | 1UUD | 2KXM | 4ZC7 | 6YL5 | 5KPY | 8HB3 | 5FKE | 8U5P |
| 3Q50 | 8I44 | 5Z1H | 7OA3 | 4E8V | 8GXB | 7EOO | 5FK3 | 4E8N | 6HBT | 6UC9 | 1LC4 |
| 4NYB | 8U60 | 1ZZ5 | 4B5R | 5FK1 | 2CKY | 5ZEJ | 8D5L | 2ET4 | 8G7E | 7OAX | 7A3Y |
| 5FK4 | 6T3N | 5LWJ | 1MWL | 3BNR | 8G4W | 2MXS | 6TF2 | 4FAX | 7E9E | 6DMC | 7D7W |
| 6YML | 6V9B | 7KVV | 4Y1J | 3GX3 | 4GPX | 3S4P | 6FZ0 | 2GDI | 6PQ7 | 7DWH | 6UP0 |
| 4E8R | 4P20 | 6YMI | 3IRW | 7TD7 | 6LAU | 7EOJ | 6T3K | 1FUF | 4FB0 | 3GX6 | 4F8U |
| 6C63 | 8I3Z | 1UTS | 6DLS | 6DMD | 3GX7 | 4P5J | 4LX5 | 7EAF | 3GES | 8EYW | 6UEJ |
| 6TFE | 6CC3 | 2PWT | 3K1V | 6E1V | 8HB8 | 2HOL | 4K31 | 4FAU | 4K32 | 6E1U | 1XPF |
| 7TDB | 4E8P | 3WRU | 1EVV | 4E8T | 4NYG | 3D2G | 4FAW | 7OAW | 3MXH | 7EOG | 6CHR |
| 1NTB | 2O3Y | 6TFH | 7TZR | 5VJ9 | 7E9I | 6XB7 | 4GPY | 2YDH | 1KOD | 7EOK | 4R0D |
| 7EOL | 6JBG | 8GXC | 6VA2 | 2ESI | 4L81 | 6VA3 | 6E1W | 7EOH | 3D2V | 2BEE | 6TF3 |
| 3SLM | 8I45 | 4OQU | 6YLB | 4F8V | 2F4U | 6QIV | 7L0Z | 4Y1M | 3GER | 3GOG | 8CF2 |
| 3E5E | 6HMO | 2ET8 | 6XRQ | 8FZA | 3D2X | 4KZD | 4FAR | 6JBF | 6LAS | 1AJU | 2HOM |
| 6C65 | 8I43 | 7EOI | 3SKL | 5V1L | 2O3W | 3DS7 | 3MUV | 7WIF | 4E8M | 5VJB | 2F4T |
| 7EDL | 3TZR | 1I9V | 2TRA | 4P3S | 5BJO | 1YRJ | 4NFO | 1F27 | 7WIB | 4NYA | 6TB7 |
| 6TF1 | 2KGP | 5V3F | 2OE5 | 7EOM | 8I46 | 6TFF | 1LVJ | 5BJP | 4Q9Q | 2O3X | 6VUI |
| 2JUK | 6E81 | 6V9D | 6T3S | 1O15 | 4AOB | 1AM0 | 3MUT | 6CK5 | 2QWY | 8HB1 | 4E8K |
| 8D28 | 4YB1 | 5T83 | 8F4O | 5FKH | 4PDQ | 1EI2 | 1Q8N | 8R8P | 6E8U | 5XZ1 | 2L94 |
| 7D82 | 6JJI | 3E5C | 7TDC | 7TZT | 6E84 | 7D81 | 6YMK | 3IQN | | | |

| **Validation Set** | | | | | | | | | | | |
|---|---|---|---|---|---|---|---|---|---|---|---|
| 3F2X | 6WTL | 6WTR | 3F2Y | 4QK8 | 6N5K | 3F2W | 4W90 | 3F2T | 6N5O | 3F4E | 6N5Q |
| 3F4G | 3F2Q | 4QKA | 6DN3 | 3F4H | 6N5S | 4QLN | 5KX9 | 4QLM | 4W92 | 6N5P | 6N5N |
| 6BFB | 6DN1 | 6DN2 | 6N5T | 3F30 | 5C45 | 4QK9 | 2YIE | | | | |

| **Test Set** | | | | | | | | | | | |
|---|---|---|---|---|---|---|---|---|---|---|---|
| 1RAW | 3DIR | 4XW7 | 3SD3 | 1YKV | 6OD9 | 5BTP | 7YC8 | 6GZR | 7ELP | 4ZNP | 6WZS |
| 3G96 | 3DIQ | 3DIL | 3DIG | 2NZ4 | 8I7N | 3B4A | 6XKO | 3L3C | 6XKN | 8UIW | 7YCG |
| 6GYV | 3G8T | 3SUX | 4LVV | 2HO7 | 3DIO | 7YCI | 6G7Z | 3G9C | 3DIM | 3MIJ | 2AU4 |
| 4QVI | 2MIY | 3SUH | 7YCH | 6Q57 | 1YLS | 4LVX | 4LVY | 3Q3Z | | | |

| **Supplementary Table 3 \| Per-target performance of SMARTPocket on the HARIBOSS test set.** | | | | | |
|---|---|---|---|---|---|
| **PDB ID** | **AUROC** | **MCC** | **F1** | **Precision** | **Recall** |
| 1RAW | 0.996 | 0.931 | 0.947 | 0.900 | 1.000 |
| 1YKV | 1.000 | 1.000 | 1.000 | 1.000 | 1.000 |
| 1YLS | 0.972 | 0.779 | 0.811 | 0.714 | 0.938 |
| 2AU4 | 0.997 | 0.858 | 0.889 | 0.889 | 0.889 |
| 2HO7 | 0.967 | 0.388 | 0.353 | 0.222 | 0.857 |
| 2MIY | 0.997 | 0.658 | 0.667 | 0.500 | 1.000 |
| 2NZ4 | 0.960 | 0.458 | 0.429 | 0.286 | 0.857 |
| 3B4A | 0.929 | 0.353 | 0.316 | 0.194 | 0.857 |
| 3DIG | 0.998 | 0.646 | 0.615 | 0.444 | 1.000 |
| 3DIL | 0.997 | 0.646 | 0.615 | 0.444 | 1.000 |
| 3DIM | 0.993 | 0.627 | 0.593 | 0.421 | 1.000 |
| 3DIO | 1.000 | 0.742 | 0.727 | 0.571 | 1.000 |
| 3DIQ | 0.928 | 0.681 | 0.696 | 0.615 | 0.800 |
| 3DIR | 0.998 | 0.646 | 0.615 | 0.444 | 1.000 |
| 3G8T | 0.936 | 0.433 | 0.400 | 0.261 | 0.857 |
| 3G96 | 0.901 | 0.433 | 0.400 | 0.261 | 0.857 |
| 3G9C | 0.920 | 0.327 | 0.286 | 0.171 | 0.857 |
| 3L3C | 0.963 | 0.382 | 0.343 | 0.214 | 0.857 |
| 3MIJ | 0.971 | 0.837 | 0.889 | 1.000 | 0.800 |
| 3Q3Z | 0.861 | 0.734 | 0.762 | 0.889 | 0.667 |
| 3SD3 | 1.000 | 1.000 | 1.000 | 1.000 | 1.000 |
| 3SUH | 0.969 | 0.594 | 0.556 | 0.385 | 1.000 |
| 3SUX | 0.956 | 0.437 | 0.421 | 0.286 | 0.800 |
| 4LVV | 1.000 | 1.000 | 1.000 | 1.000 | 1.000 |
| 4LVX | 0.997 | 0.920 | 0.933 | 0.933 | 0.933 |
| 4LVY | 0.990 | 0.733 | 0.769 | 0.833 | 0.714 |
| 4QVI | 0.629 | 0.000 | 0.000 | 0.000 | 0.000 |
| 4XW7 | 0.675 | 0.000 | 0.000 | 0.000 | 0.000 |
| 4ZNP | 0.910 | 0.869 | 0.875 | 1.000 | 0.778 |

| | | | | | |
|---|---|---|---|---|---|
| 5BTP | 0.968 | 0.798 | 0.824 | 0.778 | 0.875 |
| 6G7Z | 0.880 | 0.298 | 0.316 | 0.200 | 0.750 |
| 6GYV | 0.625 | -0.060 | 0.000 | 0.000 | 0.000 |
| 6GZR | 1.000 | 0.845 | 0.857 | 1.000 | 0.750 |
| 6OD9 | 0.924 | 0.857 | 0.875 | 0.875 | 0.875 |
| 6Q57 | 0.999 | 0.957 | 0.963 | 0.929 | 1.000 |
| 6WZS | 0.917 | 0.635 | 0.667 | 0.546 | 0.857 |
| 6XKN | 1.000 | 0.557 | 0.522 | 0.353 | 1.000 |
| 6XKO | 1.000 | 0.473 | 0.429 | 0.273 | 1.000 |
| 7ELP | 0.743 | -0.093 | 0.000 | 0.000 | 0.000 |
| 7YC8 | 0.852 | 0.159 | 0.105 | 0.057 | 0.667 |
| 7YCG | 0.668 | 0.023 | 0.044 | 0.024 | 0.286 |
| 7YCH | 0.772 | 0.078 | 0.089 | 0.054 | 0.250 |
| 7YCI | 0.754 | 0.157 | 0.140 | 0.083 | 0.429 |
| 8I7N | 0.824 | 0.095 | 0.095 | 0.057 | 0.286 |
| 8UIW | 1.000 | 0.784 | 0.778 | 0.636 | 1.000 |

| Supplementary Table 4 \| Per-target performance of SMARTPocket on the filtered Jiang test set. | | | | | |
|---|---|---|---|---|---|
| **PDB ID** | **AUROC** | **MCC** | **F1** | **Precision** | **Recall** |
| 4LVW | 0.9919 | 0.9099 | 0.9231 | 0.9231 | 0.9231 |
| 4LVZ | 1 | 1 | 1 | 1 | 1 |
| 6IZP | 0.9464 | 0.7346 | 0.8 | 0.6667 | 1 |
| 6QIQ | 1 | 1 | 1 | 1 | 1 |
| 6QIT | 0.9292 | 0.6831 | 0.8235 | 1 | 0.7 |
| 7V9E | 0.9789 | 0.6968 | 0.7 | 0.5385 | 1 |

| **Supplementary Table 5 \| Per-target performance of SMARTPocket on the time-dependent test set.** | | | | | |
|---|---|---|---|---|---|
| **PDB ID** | **AUROC** | **MCC** | **F1** | **Precision** | **Recall** |
| 7Q7Z | 0.9727 | 0.7459 | 0.75 | 0.6 | 1 |
| 8HZE | 0.9987 | 0.8828 | 0.9032 | 0.8235 | 1 |
| 8SYK | 0.9868 | 0.542 | 0.5 | 0.3333 | 1 |
| 8T5O | 0.9575 | 0.6133 | 0.5926 | 0.4211 | 1 |
| 8VPV | 1 | 0.3423 | 0.2778 | 0.1613 | 1 |
| 8ZAU | 0.9631 | 0.7511 | 0.7778 | 0.7 | 0.875 |
| 9BZ1 | 0.702 | 0.1789 | 0.2632 | 0.2174 | 0.3333 |
| 9BZC | 0.9066 | 0.465 | 0.5 | 0.375 | 0.75 |
| 9FN2 | 0.9621 | 0.6874 | 0.7 | 0.5385 | 1 |
| 9MQU | 0.8772 | 0.235 | 0.2222 | 0.1429 | 0.5 |

| **Supplementary Table 6 \| Performance of the SMARTPocket across 15 cobalamin riboswitch structures[9,10].** | | | | |
|---|---|---|---|---|
| **PDB ID** | **AUROC** | **MCC** | **Precision** | **Recall** |
| 4FRG (env8–OHCbl complex) | 0.980 | 0.692 | 0.600 | 0.900 |
| 9MFH (env2 apo) | 0.826 | 0.570 | 0.615 | 0.667 |
| 9E5H (env2- CNCbl) | 0.981 | 0.691 | 0.667 | 0.833 |
| 9E5I (env2–Cbl 4) | 0.974 | 0.847 | 0.909 | 0.833 |
| 9E5J (env2–Cbl 5) | 0.958 | 0.792 | 0.900 | 0.750 |
| 9E5K (env2–Cbl 13) | 0.868 | 0.769 | 0.909 | 0.714 |
| 9E5L (env2–Cbl 26) | 0.956 | 0.702 | 0.750 | 0.750 |
| 9E5M (env2–Cbl 29) | 0.967 | 0.744 | 0.818 | 0.750 |
| 9E5O (env2–Cbl 32) | 0.993 | 0.905 | 1.000 | 0.846 |
| 9E5P (env2–Cbl 33) | 0.980 | 0.775 | 0.786 | 0.846 |
| 9E5Q (env2–Cbl 36) | 0.980 | 0.847 | 0.909 | 0.833 |
| 9E5R (env2–Cbl 37) | 0.963 | 0.680 | 0.727 | 0.727 |
| 9E5S (env2–Cbl 63) | 0.934 | 0.546 | 0.529 | 0.750 |
| 9E5T (env2–Cbl 64) | 0.953 | 0.672 | 0.643 | 0.818 |
| 9ELR (env2–Cbl 42) | 0.976 | 0.744 | 0.818 | 0.750 |

Note: Both *env8* and *env2* are closely related cobalamin riboswitches (sharing 95% sequence identity), isolated from ocean metagenomic samples. OHCbl (hydroxocobalamin) and CNCbl (cyanocobalamin) are two widely used forms of vitamin B12. Cbl (cobalamin) followed by a number (e.g., Cbl 4, Cbl 29) denotes synthetic cobalamin derivatives with varied β-axial moieties; X-ray crystallography of these complexes revealed that bulkier β-axial groups displace nucleotide A20 from the RNA core, exposing a cryptic binding pocket. The apo structure (9MFH) is the ligand-free form of *env2*. Detailed information can be found in the corresponding references[9,10].

| **Supplementary Table 7 \| Distribution of RNA structures with varying numbers of small molecule binders in the HARIBOSS benchmark dataset.** | |
|---|---|
| **Number of ligands in the RNA structure** | **Number of samples** |
| 1 | 270 |
| 2 | 115 |
| 3 | 20 |
| 4 | 15 |
| 5 | 7 |
| 6 | 2 |
| 7 | 4 |

| **Supplementary Table 8 \| Pocket-level structural motif annotations for RNA targets in the HARIBOSS[1], filtered Jiang[2] and time-dependent tests, as retrieved from RNA 3D Hub[3].** | | | |
|---|---|---|---|
| **PDB ID** | **Motif Annotation** | **PDB ID** | **Motif Annotation** |
| 7ELP | Bulge loop | 7YC8 | Internal loop |
| 7V9E | Bulge loop | 7YCG | Internal loop |
| 7Q7Z | Bulge loop | 2AU4 | Internal loop |
| 6OD9 | Double sheared | 6QIQ | Internal loop |
| 5BTP | Double sheared | 6QIT | Internal loop |
| 6WZS | Double sheared | 8HZE | Internal loop |
| 6GYV | Internal loop | 3G96 | Junction loop |
| 6XKN | Hairpin loop | 2NZ4 | Junction loop |
| 4LVX | Hairpin loop | 3DIL | Junction loop |
| 2HO7 | Hairpin loop | 3SUX | Junction loop |
| 3SD3 | Hairpin loop | 3SUH | Junction loop |
| 3B4A | Hairpin loop | 3L3C | Junction loop |
| 4ZNP | Hairpin loop | 3DIR | Junction loop |
| 6XKO | Hairpin loop | 3DIQ | Junction loop |
| 4LVY | Hairpin loop | 3G8T | Junction loop |
| 4LVV | Hairpin loop | 3DIO | Junction loop |
| 4LVW | Hairpin loop | 6GZR | Junction loop |
| 4LVZ | Hairpin loop | 6Q57 | Junction loop |
| 6IZP | Internal loop | 3DIG | Junction loop |
| 8VPV | Hairpin loop | 3DIM | Junction loop |
| 8T5O | Hairpin loop | 3G9C | Junction loop |
| 9MQU | Hairpin loop | 9FN2 | Junction loop |
| 9BZ1 | Hairpin loop | 8ZAU | Junction loop |
| 9BZC | Hairpin loop | 3Q3Z | Internal loop |
| 8SYK | Hairpin loop | 7YCH | Internal loop |
| 1RAW | Internal loop | 1YKV | Internal loop |
| 4QVI | Internal loop | 1YLS | Internal loop |
| 7YCI | Internal loop | 2MIY | Pseudoknot |
| 4XW7 | Internal loop | 8UIW | Hairpin loop |
| 8I7N | Internal loop | 6G7Z | Junction loop |

| **Supplementary Table 9 \| Per PDB performance of the SMARTPocket on apo-holo pairs** | | | | | |
|---|---|---|---|---|---|
| **PDB** | **AUROC** | **MCC** | **F1** | **Precision** | **Recall** |
| 3SUX | 0.9916 | 0.8898 | 0.8889 | 1 | 0.8 |
| 3SUY | 0.9926 | 0.7895 | 0.8 | 0.8 | 0.8 |
| 3D0U | 1 | 1 | 1 | 1 | 1 |
| 3D0X | 1 | 1 | 1 | 1 | 1 |
| 2YIE | 0.9909 | 0.7906 | 0.8 | 0.6667 | 1 |
| 6WJR | 0.7474 | 0.2517 | 0.3448 | 0.2941 | 0.4167 |
| 4LVX | 0.9788 | 0.9186 | 0.9286 | 1 | 0.8667 |
| 7KD1 | 0.7392 | 0.4818 | 0.4211 | 1 | 0.2667 |
| 6XJQ | 0.9776 | 0.6893 | 0.7 | 0.5385 | 1 |
| 6XJZ | 0.9944 | 0.7658 | 0.7778 | 0.6364 | 1 |
| 2L8H | 0.9737 | 0.7162 | 0.8182 | 0.75 | 0.9 |
| 1ANR | 0.3368 | -0.2213 | 0.3077 | 0.25 | 0.4 |

### SUPPLEMENTARY NOTES

#### Supplementary Note 1 | Multiple chains and ligands

A quantitative analysis of the HARIBOSS[1] benchmark dataset, comprising 433 RNA structures, reveals that single-chain RNAs predominate, accounting for 301 structures (Supplementary Table 7). The remaining 132 structures contain multiple RNA chains. Among these, two-chain complexes are the most frequent (n = 119), followed by three-chain (n = 9) and four-chain (n = 4) structures. This distribution indicates that nearly one-third of RNA-ligand complexes involve multiple RNA chains, highlighting the importance of accurately capturing multiple chain interacting with ligand in predictive modeling.

In terms of ligand associations, most RNAs bind a single ligand (270 cases). A total of 163 RNAs interact with multiple ligands, with two-ligand complexes being the most common (n = 115). A few structures exhibit extensive ligand binding, accommodating up to seven ligands.

### Supplementary Note 2 | Docking

#### RNA–ligand structure preparation and ligand quantum mechanical optimization

RNA–ligand complexes were processed using an automated structural preparation workflow implemented in Python. The workflow accepts experimentally determined structures in either Protein Data Bank (PDB) or Macromolecular Crystallographic Information File (mmCIF/CIF) format as input. Structural parsing and manipulation were performed using Biopython[11] and RDKit-based[12] utilities.

#### Extraction of RNA and small molecule components

For each input structure, all solvent molecules, crystallographic waters, ions, buffer components, and protein chains were removed. Only RNA residues and bound small molecule ligands were retained for downstream analysis. Ligands were identified based on residue annotations while excluding canonical RNA nucleotides, modified nucleotides integrated into the RNA backbone, water molecules, and common crystallographic ions.

The RNA and ligand coordinates were subsequently separated into independent structural files. RNA coordinates were saved in PDB format for docking and structural analyses. Small molecule ligands were exported in both PDB and Structure Data File (SDF) formats to preserve bond order and chemical topology information required for quantum mechanical optimization and ligand preparation.

#### Ligand structural validation and missing atom correction

Prior to ligand optimization, extracted ligands were validated against their corresponding reference chemical structures using SMILES-based pattern matching. Ligands exhibiting missing heavy atoms, incomplete connectivity, incorrect bond orders, or valence inconsistencies relative to the reference structure were automatically identified and corrected before downstream processing. Ligands with incomplete structural definitions were corrected where possible using RDKit chemical sanitization procedures and template-based bond reconstruction[12].

### Quantum mechanical geometry optimization

Ligand geometries were optimized using the Psi4 quantum chemistry package[13]. Initial three-dimensional coordinates derived from the experimental structures were subjected to quantum mechanical geometry refinement using a user-defined basis set and level of theory. Typical calculations employed Hartree–Fock (HF)[14] or density functional theory (DFT)-based[15] optimization protocols depending on user selection.

Geometry optimization was performed iteratively until convergence criteria for energy, gradient, and atomic displacement were satisfied. For Psi4 optimizations, convergence was defined according to the program's default geometry optimization thresholds, including maximum force ($< 4.5 \times 10^{-4}$ Hartree Bohr$^{-1}$), RMS force ($< 3.0 \times 10^{-4}$ Hartree Bohr$^{-1}$), maximum displacement ($< 1.8 \times 10^{-3}$ Bohr), and RMS displacement ($< 1.2 \times 10^{-3}$ Bohr). Following each optimization cycle, convergence status was automatically evaluated by parsing Psi4 output files. Structures failing to converge within the default optimization cycle limit were automatically resubmitted with an increased maximum number of optimization steps until convergence was achieved or the structure was flagged for manual inspection. This iterative optimization strategy ensured stable and chemically realistic ligand geometries prior to docking preparation.

### Ligand Protonation State Assignment

Optimized ligands were protonated at physiological pH (7.4) using the LigPrep module from the Schrödinger software suite. Protonation states, tautomeric forms, and stereochemical assignments were generated under Epik-based ionization state prediction settings corresponding to near-physiological conditions[16]. The resulting protonated structures were subsequently used for charge derivation and docking preparation.

**RESP Partial Charge Assignment**

Electrostatic partial charges for each ligand were assigned using the Restrained Electrostatic Potential (RESP) methodology[17]. Electrostatic potentials were calculated from the quantum mechanically optimized ligand geometries, and RESP fitting was performed to derive atom-centered partial charges suitable for molecular docking and molecular mechanics simulations. The RESP charge derivation procedure ensured physically consistent electrostatic representations of ligand molecules and improved compatibility with downstream docking and free-energy calculations.

**Identification of the best docking pose**

Following docking, all generated poses were first ranked according to their predicted binding free energy as reported by the docking scoring function. Because docking scoring functions typically exhibit uncertainties on the order of 1–2 kcal/mol and often cannot reliably discriminate among closely related binding modes, poses with docking scores within 2 kcal/mol of the top-ranked pose were considered energetically equivalent and retained for further analysis[18,19]. This energy window is widely used in docking studies because multiple binding conformations separated by less than approximately 2 kcal/mol are expected to be thermally accessible at physiological temperatures and frequently represent alternative local minima within the same binding pocket[19]. Each retained pose was subsequently aligned to the experimentally determined RNA–ligand complex, and the heavy-atom root-mean-square deviation (RMSD) relative to the native ligand conformation was calculated. Among all retained poses, the pose exhibiting the lowest RMSD to the experimentally observed ligand was designated as the best recovered pose. This approach, also used in other published reports[19], minimizes artifacts arising from small scoring-function errors while providing a robust assessment of whether the docking protocol successfully sampled a near-native binding mode.

Docking calculations were conducted using the AutoDock4 empirical free-energy scoring function, which evaluates intermolecular interactions through van der Waals, hydrogen-bonding, electrostatic, desolvation, and torsional energy terms[20]. The Solis–Wets local search algorithm was employed for conformational optimization[20]. For each RNA–ligand pair, 20 independent docking poses were generated and ranked according to their predicted binding free energies. Two docking strategies were evaluated: blind docking and SMARTPocket-guided docking.

For blind docking, the search box was defined to encompass the entire RNA structure. The dimensions of the docking box were automatically determined from the spatial extent of the RNA target by calculating the minimum and maximum atomic coordinates along each Cartesian axis and expanding the box to fully cover the complete RNA molecule[20]. This approach ensured that all solvent-accessible surfaces and potential ligand-binding regions of the RNA could be explored during docking.

For SMARTPocket-guided docking, nucleotide-level pocket predictions generated by SMARTPocket were used to define the docking region. Specifically, the nucleotides identified as belonging to the predicted binding pocket were selected, and a docking box was constructed to encompass all atoms within these residues. To ensure unrestricted ligand sampling within the predicted pocket, the search box was further expanded by a distance proportional to the maximum molecular dimensions of the ligand. This additional padding allowed complete translational and rotational freedom of the ligand within the search region while maintaining a focused exploration of the SMARTPocket-predicted binding site. Consequently, blind docking sampled the entire RNA surface, whereas SMARTPocket-guided docking restricted sampling to a biologically relevant region while preserving sufficient conformational space for ligand movement and pose optimization.
